# Disease Resistance Genetics and Genomics in Octoploid Strawberry

**DOI:** 10.1101/646000

**Authors:** Christopher Barbey, Seonghee Lee, Sujeet Verma, Kevin A. Bird, Alan E. Yocca, Patrick P. Edger, Steven J. Knapp, Vance M. Whitaker, Kevin M. Folta

## Abstract

Octoploid strawberry (*Fragaria × ananassa*) is a valuable specialty crop, but profitable production and availability are threatened by many pathogens. Efforts to identify and introgress useful disease resistance genes (R-genes) in breeding programs are complicated by strawberry’s complex octoploid genome. Recently-developed resources in strawberry, including a complete octoploid reference genome and high-resolution octoploid genotyping, enable new analyses in strawberry disease resistance genetics. This study characterizes the complete R-gene collection in the genomes of commercial octoploid strawberry and two diploid ancestral relatives, and introduces several new technological and data resources for strawberry disease resistance research. These include octoploid R-gene transcription profiling, *d*N/*d*S analysis, eQTL analysis and RenSeq analysis in cultivars. Octoploid fruit transcript expression quantitative trait loci (eQTL) were identified for 77 putative R-genes. R-genes from the ancestral diploids *Fragaria vesca* and *Fragaria iinumae* were compared, revealing differential inheritance and retention of various octoploid R-gene subtypes. The mode and magnitude of natural selection of individual *F. × ananassa* R-genes was also determined via *d*N/*d*S analysis. R-gene sequencing using enriched libraries (RenSeq) has been used recently for R-gene discovery in many crops, however this technique somewhat relies upon *a priori* knowledge of desired sequences. An octoploid strawberry capture-probe panel, derived from the results of this study, is validated in a RenSeq experiment and is presented for community use. These results give unprecedented insight into crop disease resistance genetics, and represent an advance towards exploiting variation for strawberry cultivar improvement.

## INTRODUCTION

Cultivated strawberry (*Fragaria ×ananassa*) is an important specialty crop that is cultivated world-wide for its sweet and flavorful fruit. However, marketable yields and post-harvest quality are significantly affected by disease. The strawberry fruit presents a vulnerable target for microbial pathogens (Farzaneh *et al.*, 2015), as it is soft, moist, carbohydrate rich, and subject to damage from forces as seemingly innocuous as rain (Herrington *et al.*, 2011). Genetic disease resistance has been a long-standing breeding priority. While breeders have made progress in producing varieties with tolerance to some pathogens, growers remain dependent on exogenous crop protection strategies to reduce pathogen loads (Cordova *et al.*, 2017).

Plant R-genes are mediators of resistance to specific pathogens via effector triggered immunity, which results in the hypersensitive response and cell death (Amil-Ruiz *et al.*, 2018). R-genes require a high degree of regulation to maintain homeostatic transcript levels to mitigate off-target protein interactions (Hammond-Kosack and Jones, 1997). For this reason, many classes of functional R-genes are expressed at low levels unless elicited by pathogens (Lai and Eulgem, 2017), contributing to the challenges of R-gene genomic and functional annotation. About 60% of characterized plant R-genes contain nucleotide-binding (NB-ARC) and leucine-rich-repeat (LRR) domains, and are referred to NLR genes (Funk *et al.*, 2018). Plant R-genes are frequent targets for genetic improvement via breeding and genetic engineering (Djian-Caporalino *et al.*, 2014, Baumgartner *et al.*, 2015), and gene editing methods may accelerate their introduction into already-elite varieties. However, progress has been hindered because relatively few R-genes conferring novel resistance have been characterized (Amil-Ruiz *et al.*, 2018). This problem is appreciable in strawberry, where the genetic complexity of octoploid cultivars presents unique challenges for functional identification and cloning of causal variants. An analysis of diploid R-genes across the Rosaceae genus was previously conducted (Arya *et al.*, 2014). New genetic resources for high-resolution genotyping in octoploid strawberry have resulted in the recent identification of several disease resistance loci (Roach *et al.*, 2016, Mangandi *et al.*, 2017, Anciro *et al.*, 2018, Pincot *et al.*, 2018, Salinas *et al.*, 2018, Verma *et al.*, 2018). However, the specific genes mediating resistance in these QTL intervals typically remain unresolved, as genomic resources for octoploid strawberry have not kept pace with genetic mapping.

Cultivated strawberry shares common ancestors with the extant diploid species *F. vesca, F. iinumae, F. nipponica*, and *F. viridis* (Edger *et al.*, 2019). A high-quality octoploid strawberry genome has been recently developed (Edger *et al.*, 2019), enabling new kinds analyses and improved resolution compared with previous studies involving *Fragaria* NLRs (Jia *et al.*, 2015, Zhong *et al.*, 2018). Analysis of this *F. ×ananassa* ‘Camarosa’ genome identified the repertoire of octoploid R-gene sequences and further demonstrated a general genomic retention bias towards *F. vesca*-like sequences (Edger *et al.*, 2019).

This research compares R-genes from octoploid strawberry with its diploid ancestors and provides additional analysis into the genetic control of R-gene expression and retention patterns. Additional bias towards retention of *F. vesca-*like R-genes was detected in octoploid strawberry, beyond the bias observed in non-R-gene coding sequences. This finding provides insight into potential practical drivers of biased gene retention. Conserved domains were compared to describe specific R-gene phylogenic relationships. The octoploid genome was used to assemble 61 fruit transcriptomes, and used to discover subgenomic expression quantitative trait loci (eQTL) for R-genes expressed in octoploid fruit. Data from the octoploid ‘Camarosa’ strawberry gene expression atlas (Sánchez-Sevilla et al., 2017) was also used to determine R-gene transcript accumulation throughout the strawberry plant.

Resistance gene enrichment and sequencing (RenSeq) is an advantageous method for sequencing R-genes (Andolfo *et al.*, 2014), and is likely to be very useful for *de novo* resolution of causal mutations (Witek *et al.*, 2016). This method can be used to identify casual mutations within existing disease resistance QTL. For this purpose, a novel octoploid strawberry RenSeq capture probe library was designed using the R-genes identified in this analysis. This panel was experimentally validated using the University of Florida breeding germplasm. The results demonstrate robust capture and resequencing of octoploid and diploid R-genes using only short second-generation sequence reads and with relatively deep genomic multiplexing.

This report characterizes the complete R-gene collection in the genomes of commercial octoploid strawberry and two diploid ancestral relatives, providing the genome-level resolution necessary for fully exploiting genetic disease resistance in strawberry. This research introduces several new technology and data resources that now may be applied in study of strawberry disease resistance.

## RESULTS

### Octoploid and Diploid R-genes

The genomes of octoploid ‘Camarosa’, diploid *F. vesca*, and diploid *F. iinumae* were analyzed for R-gene signatures. The *F.iinumae* genome was selected to represent the closely-related ‘old world’ diploid ancestors *F.iinumae, F. nipponica* and *F. viridis*, which each have highly similar but fragmented genomic assemblies.

Putative R-genes were identified based on protein domain and motif analysis, which identified gene models with traditional NLR-type domains, including coiled coil (CC), Toll Interleukin Receptor-like (TIR), Leucine Rich Repeat (LRR), and Nucleotide Binding - APAF-1 (apoptotic protease-activating factor-1), R proteins and CED-4 (*Caenorhabditis elegans* death-4 protein) (NB-ARC). Gene models with NLR-type domains that are not highly specific to NLR sequences (e.g. LRR domains) were included if there was also supporting evidence of an additional NLR-associated motif. BLAST2GO annotated disease resistance associated genes not meeting these criteria were analyzed manually, leading to the intentional inclusion of many putative Receptor-like Kinase (RLK-type) R-genes in this analysis.

Octoploid *F. ×ananassa* ‘Camarosa’ carries 1,962 putative resistance genes (1.82% of all genes), including 975 complete or truncated NLR genes (Table 1). NLR gene content is similar in genic proportion to the 367 complete or truncated NLR genes in *F. vesca* (1.09% of all genes) and 387 in *F. iinumae* (0.5% of all genes) (Table 1). Traditional NLR domains including Coiled Coil (CC), Toll Interleukin Receptor-like (TIR), Leucine Rich Repeat (LRR), and Nucleotide Binding - APAF-1 (apoptotic protease-activating factor-1), R proteins and CED-4 (*Caenorhabditis elegans* death-4 protein) (NB-ARC) domains comprise the majority of domain classes in all predicted resistance gene models in diploid and octoploid strawberry accessions (Figure 1). In many categories, the three genomes show somewhat dissimilar ratios of relative NLR-subtype content (Table 1). These include biases towards TIR-only proteins in *F. vesca* and CNL-type R-genes in *F. innumae*. Octoploid ‘Camarosa’ is proportionally intermediate for many NLR categories, relative to *F. vesca and F. innumae*. A high proportion of TIR-NB and TIR-NB-LRR-type NLR-genes is observed in ‘Camarosa’. However, the overall proportion of TIR-containing genes is similar. The Resistance to Powdery Mildew 8 (RPW8) domain, a disease resistance domain associated with broad-spectrum mildew resistance in Arabidopsis, appears frequently in strawberry and is present in 136 (13.9%) of octoploid NLRs (Table 1). Basic trends in NLR-subtype genomic content in ‘Camarosa’ does not more strongly resemble either *F. vesca* or *F. innumae*.

**Table 1.** NLR-gene subtype distribution across three strawberry species

|  | *cv. Camarosa* | *F. vesca* | *F. iinumae* |
| --- | --- | --- | --- |
| NB | 19.8% | 17.7% | 11.6% |
| NB-LRR | 2.3% | 1.9% | 5.2% |
| **NB-only type** | **22.1%** | **19.6%** | **16.8%** |
| CC-NB | 16.7% | 12.3% | 32.6% |
| CC-NB-LRR | 3.0% | 2.5% | 2.1% |
| **CNL-type** | **19.7%** | **14.7%** | **34.6%** |
| RPW | 9.3% | 10.9% | 5.9% |
| RPW-NB | 4.6% | 8.4% | 3.4% |
| **RPW-type** | **13.9%** | **19.3%** | **9.3%** |
| TIR | 13.7% | 37.1% | 22.2% |
| TIR-NB | 20.0% | 4.9% | 13.2% |
| TIR-NB-LRR | 10.6% | 4.4% | 3.9% |
| **TNL-type** | **44.3%** | **46.3%** | **39.3%** |

Relative content of NLR-gene subtypes and truncated subtypes are shown.

**Figure 1.**
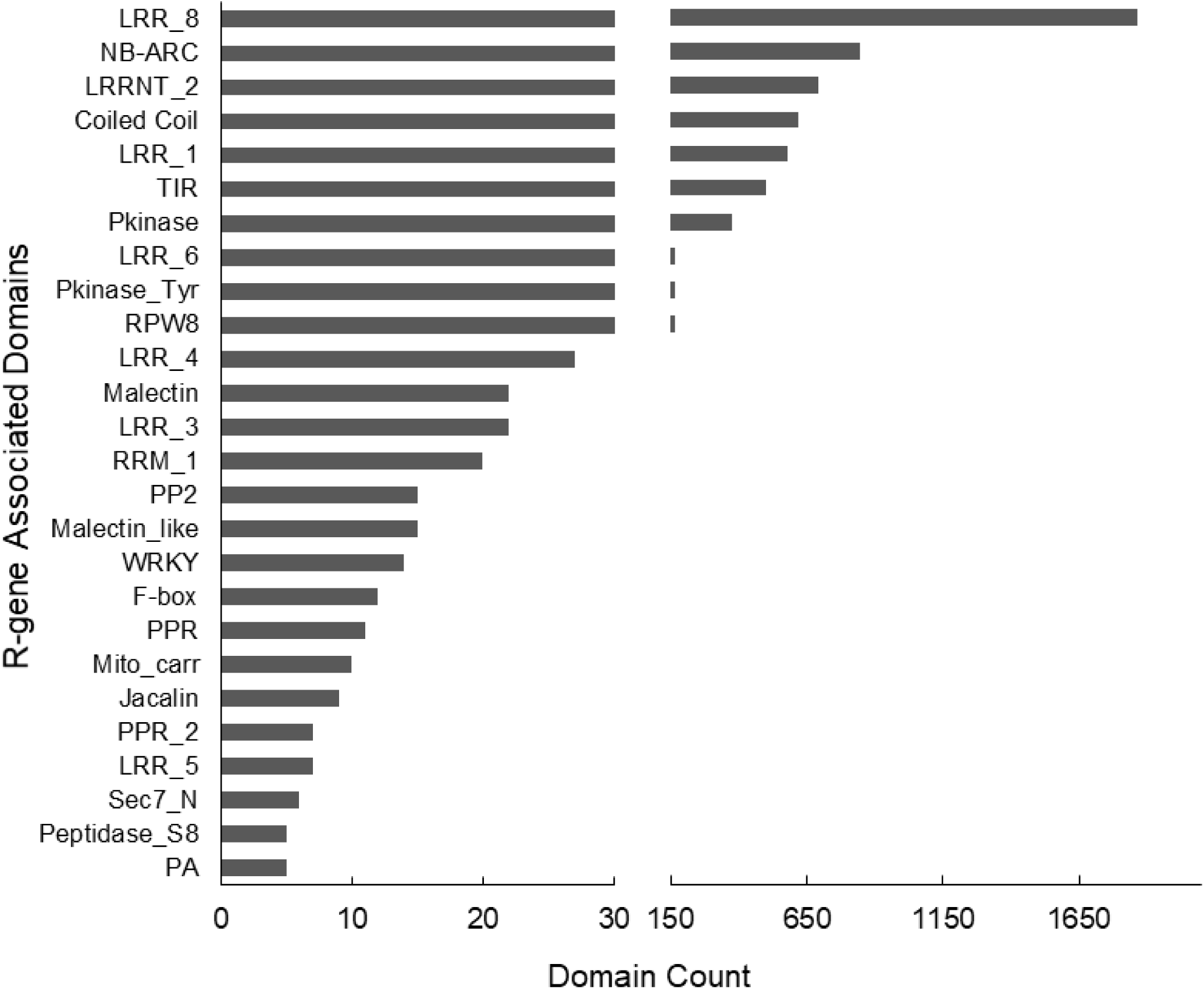
Canonical and non-Canonical Domains in Strawberry R-Genes. Classic TNL/CNL-type R-gene domains (TIR, NB-ARC, LRR, etc.) comprise the majority of domain classes in predicted R-genes, however a number of atypical domains are observed in high frequency. Domains below a count of five are not shown.

In ‘Camarosa’, 750 genes contain at least one NB-ARC domain, which is the most characteristic domain of NLR-type R-genes (Table 1). The ratio of ‘Camarosa’ NB-ARC-containing genes to total predicted gene content (1:144) is higher than in *F. vesca* (1:171) and *F. iinumae* (1:262), possibly indicating diversifying selection of NLR genes in octoploid *F. ×ananassa*. A substantial number of atypical domains are present on strawberry R-genes, including malectin-like carbohydrate-binding domains, RNA-binding domains, transcription factor-like WRKY and F-box domains, and several types of protein kinase domains (Figure 1).

Tandem clusters of R-genes were observed in all three of the analyzed strawberry genomes. The phenomena of R-gene expansion through tandem duplication is exemplified in the RPW8-containing R-gene class. Of the seventy-one RPW-containing R-genes in *F. vesca*, all but seven reside in one of a few genomic clusters (Figure S1). The major RPW cluster observed in *F. vesca* chromosome 1 is strongly retained in ‘Camarosa’ (Figure S2). Similar R-gene hotspots are observed throughout the diploid and octoploid strawberry reference genomes. Genome annotations for all NLR domains in ‘Camarosa’ and *F. vesca* genotype Hawaii 4 v2.0 (Tennessen *et al.*, 2014) are provided in File S1. Annotations are also available on the JBrowse web-based genome browser at the Genome Database for the Rosaceae (www.rosaceae.org).

### Phylogenetic Analysis of Strawberry R-genes

The conserved NB-ARC domains from ‘Camarosa’, *F. vesca*, and *F. iinumae* were compared via maximum likelihood analysis to examine evolutionary trends among NLR genes. NB-ARCs from all three genomes phylogenetically organized mostly according to their extended R-gene domain structures, with TNLs, CNLs, and RPW-associated NB-ARCs forming clades based on this criteria (Figure 2). Minor NLR subtypes, such as WRKY-associated NLR genes, also sorted into a unique subclade based only on NB-ARC sequence. Multiple distinct clades with identical domain architectures were detected, and in a few cases these subclades are relatively distant from one another.

**Figure 2.**
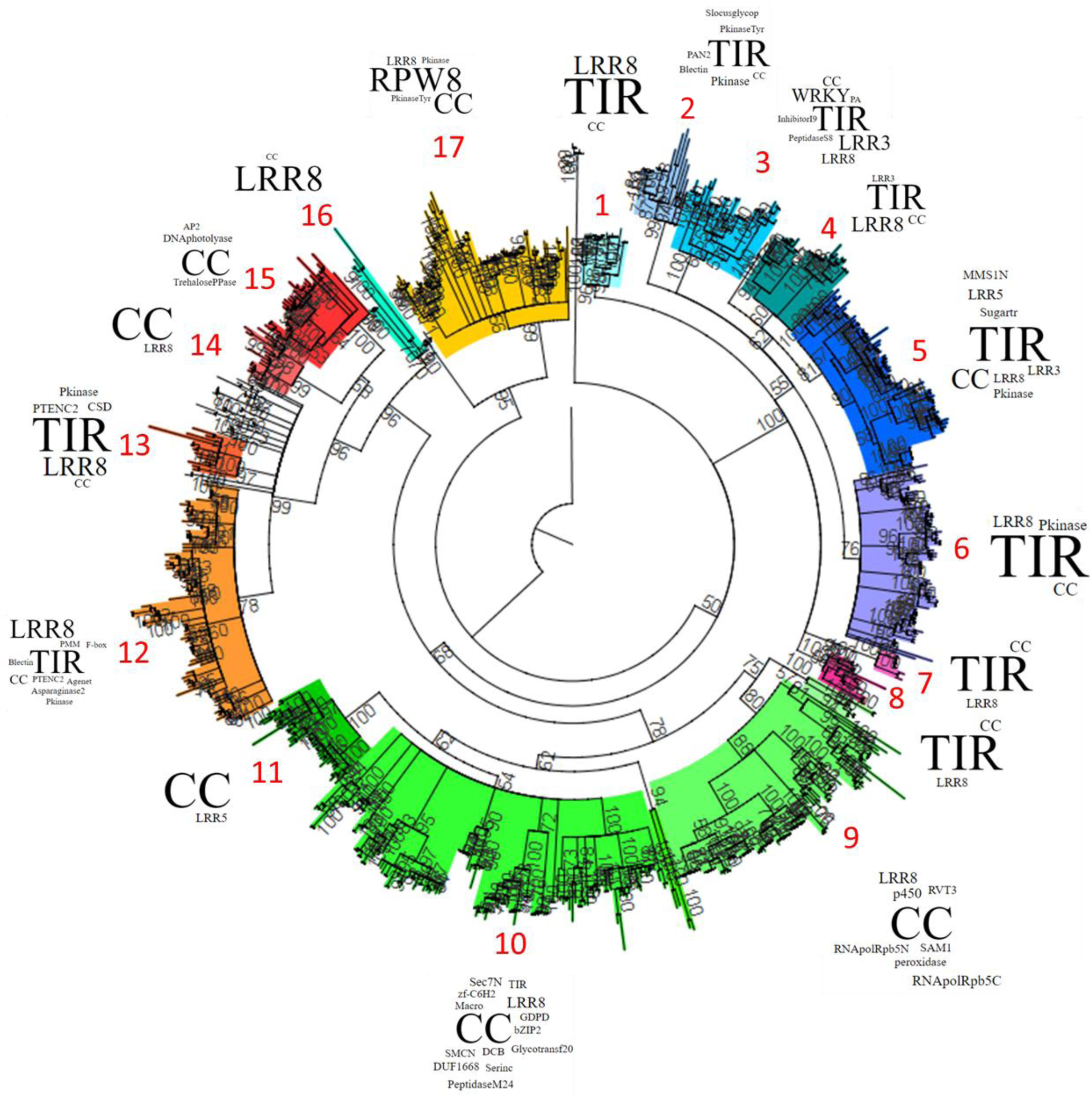
Phylogenetic Relationship of NB-ARC domains in *F. vesca, F. iinumae*, & *F. × ananassa* ‘Camarosa’. A. Full-length NB-ARC domains from strawberry *spp*. organize into clades based on NLR-gene subtype (CC, TIR, NB-ARC, LRR, and RPW-containing combinations). Maximum likelihood bootstrap values (100 replicates) above a threshold of 50% are shown with the NB-ARC domain from human *Apaf1* as the outgroup. Word sizes correspond to relative domain content within each clade.

### R-gene Transcript Accumulation

Raw RNAseq expression data from different tissues of ‘Camarosa’, derived from the octoploid strawberry gene expression atlas (Sánchez-Sevilla *et al.*, 2017), were reassembled based on the ‘Camarosa’ genome. A majority of ‘Camarosa’ NLR genes are predominantly expressed in the roots and leaves (Figure 3A-B). Comparatively few NLRs are predominantly or specifically expressed in the mature receptacle. Most NLR type R-genes are broadly specific to only one or two tissues. Expressed NLR genes from root, leaf, green and white receptacles show remarkably poor overlap. Overall NLR transcript accumulation is correlated across all stages of receptacle development, with strongest expression in the earlier stages and decreasing with maturity in both the receptacle and achene. Complete R-gene expression values for each tissue are provided in Table S1.

**Figure 3.**
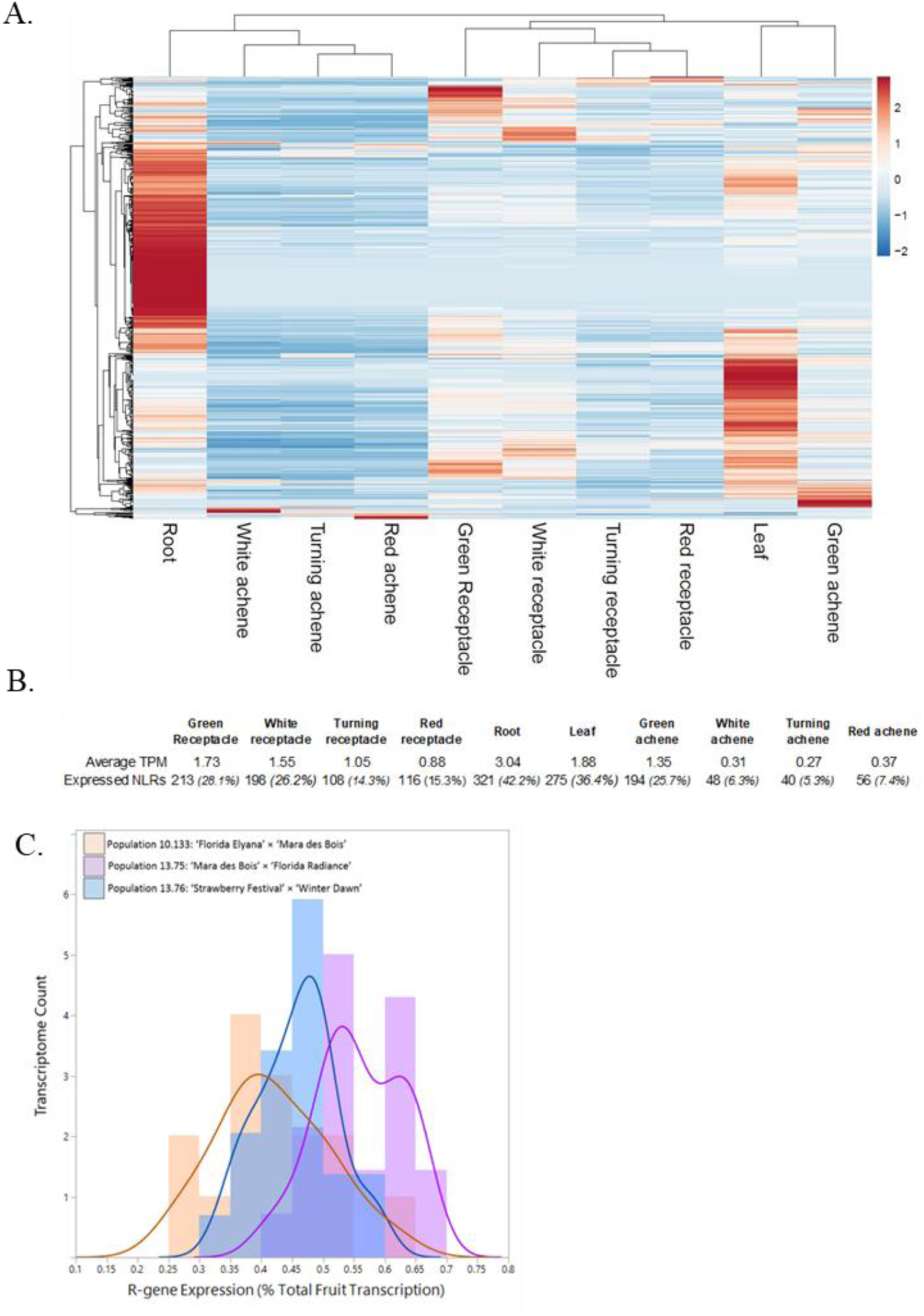
RNAseq Expression of Octoploid NLR genes. A. Tissue-based heatmap of scaled NLR transcript accumulation. B. Averaged NLR transcript abundance (TPM) and total number of putative expressed NLRs (TPM >1) are reported. The percent of all 756 NLR genes putatively expressed in each tissue are shown in parentheses. C. Generalized R-gene fruit expression across three segregating populations (n=61).

**Figure 4.**
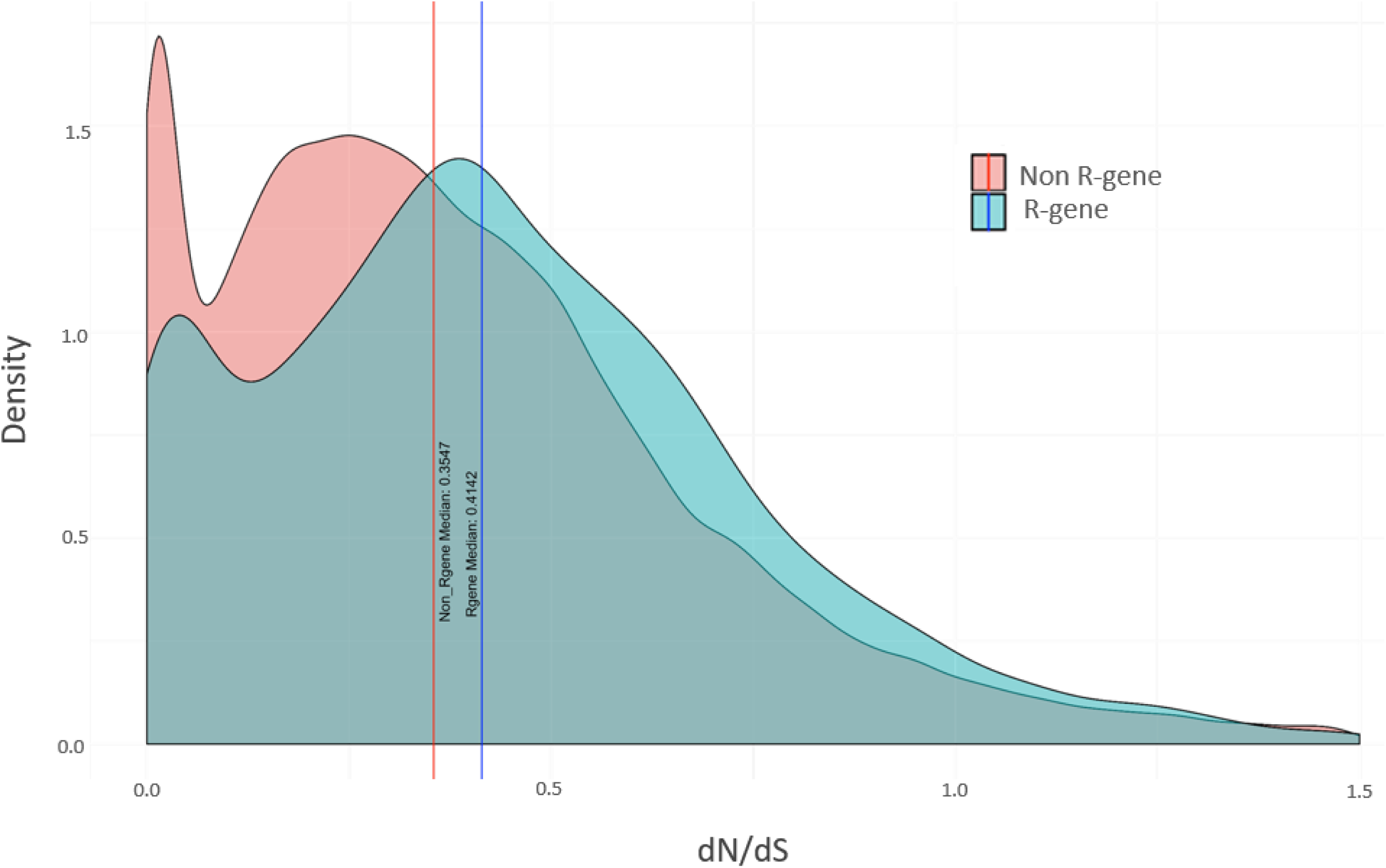
Evolutionary Pressures on *F. ×ananassa* R-genes. The median *d*N/*d*S ratio for R-genes (0.4142) is higher than for non R-genes (0.3547). Density curves for *F. ×ananassa* R-genes (blue) and non R-genes (red) are calculated based on comparison to the closest ancestral diploid homolog from *F. vesca*.

Mature receptacle transcriptomes from 61 field-grown individuals of three octoploid populations reveal broadly stable R-gene expression levels (SD 0.09). R-genes comprise 1.8% of the predicted gene models in the ‘Camarosa’ genome, but represent an average of only 0.48% of total transcripts in the mature receptacle (Figure 3C). Minute but statistically significant absolute differences were observed between each of the three populations [F (2,59) = 19.06, p < .00001]. To explore possible biases in the gene expression analysis caused by confounding environmental factors, principal component analysis (PCA) was performed on all RNAseq assemblies including two replicates of ‘Mara des Bois’ fruit harvested in different seasons (Figure S2). Total transcript-accumulation variation clusters most strongly according to familial relationship, with the ‘Mara des Bois’ replicates showing similar expression patterns. A measured amount of variation due to environmental influence can also be seen, as the two ‘Mara des Bois’ RNAseq replicates cluster somewhat more closely with their co-harvested progeny.

### R-gene eQTL in 61 Strawberry Fruit Transcriptomes

eQTL analysis was performed to evaluate heritable genotypic effects on R-gene transcription using 61 octoploid IStraw35 genotypes and mature fruit transcriptomes. This analysis identified 77 R-gene-like sequences with at least one highly-significant locus explaining differential expression (3.9% of octoploid genome-predicted R-genes). These R-genes include 53 NB-ARC containing genes, comprised of 25 TNL’s, 14 CNL’s, 3 NBS-RPW proteins, and 11 NB-ARC-only proteins (File S1). The majority of remaining eQTL genes are TIR, LRR, and RPW-only genes. As the ‘Camarosa’ genomic locus of each transcript is known, *cis* vs *trans* eQTL status was determined. Of 77 significant R-gene eQTL transcripts, all 77 R-genes are regulated via a *cis*-genetic locus (Table 2), of which 24 R-genes are also under regulation of an additional second *trans*-eQTL (Table 3). No solely *trans*-eQTL were discovered among this set of R-genes. The most significant IStraw35 SNP marker name and position for each R-gene transcript is provided with the eQTL phase, minor allele frequency, p-value (FDR-adjusted), heritability estimate, expression in parental lines, and BLAST2GO description.

**Table 2.**
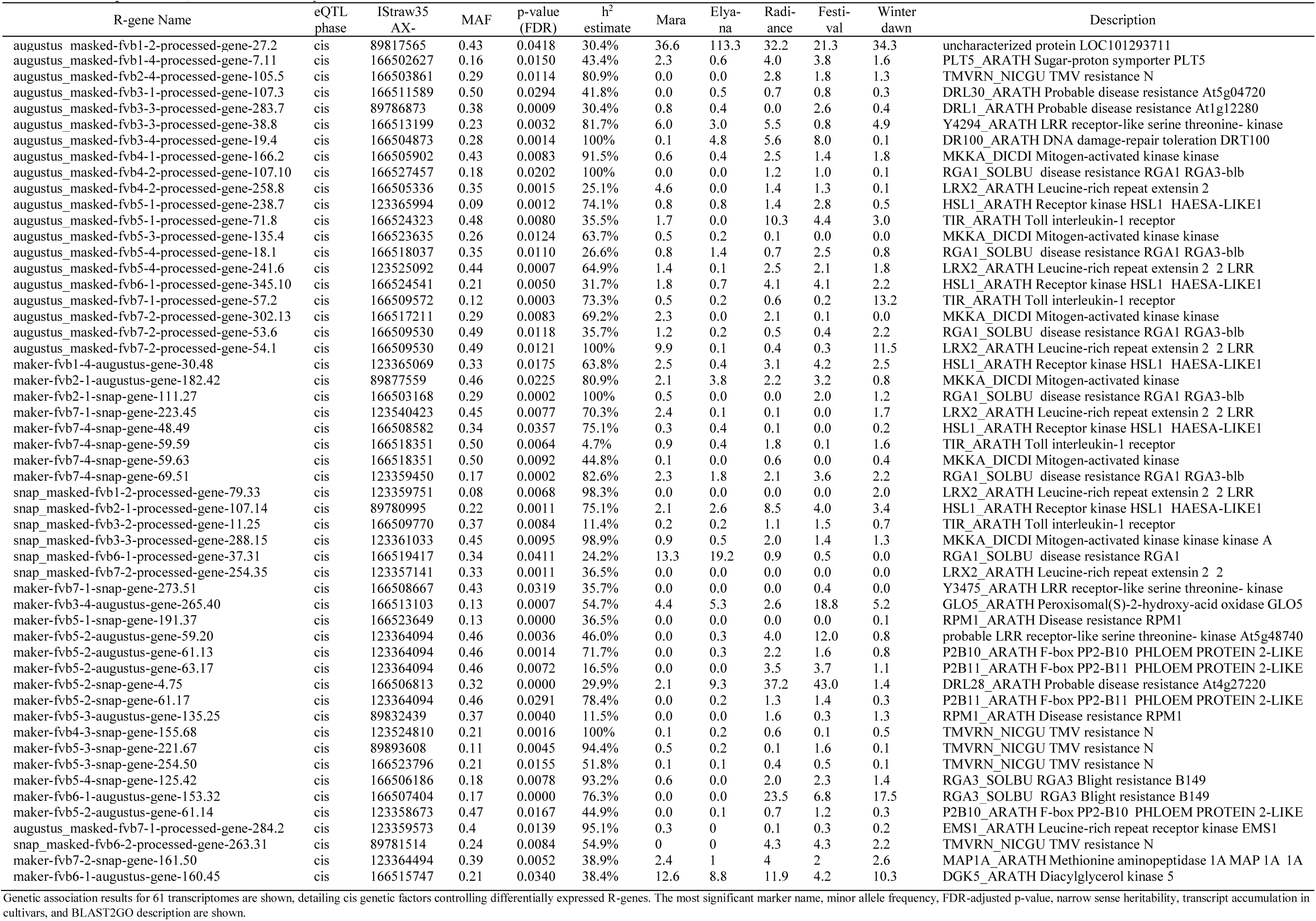
Cis Expression-QTL for Strawberry Fruit

**Table 3.**
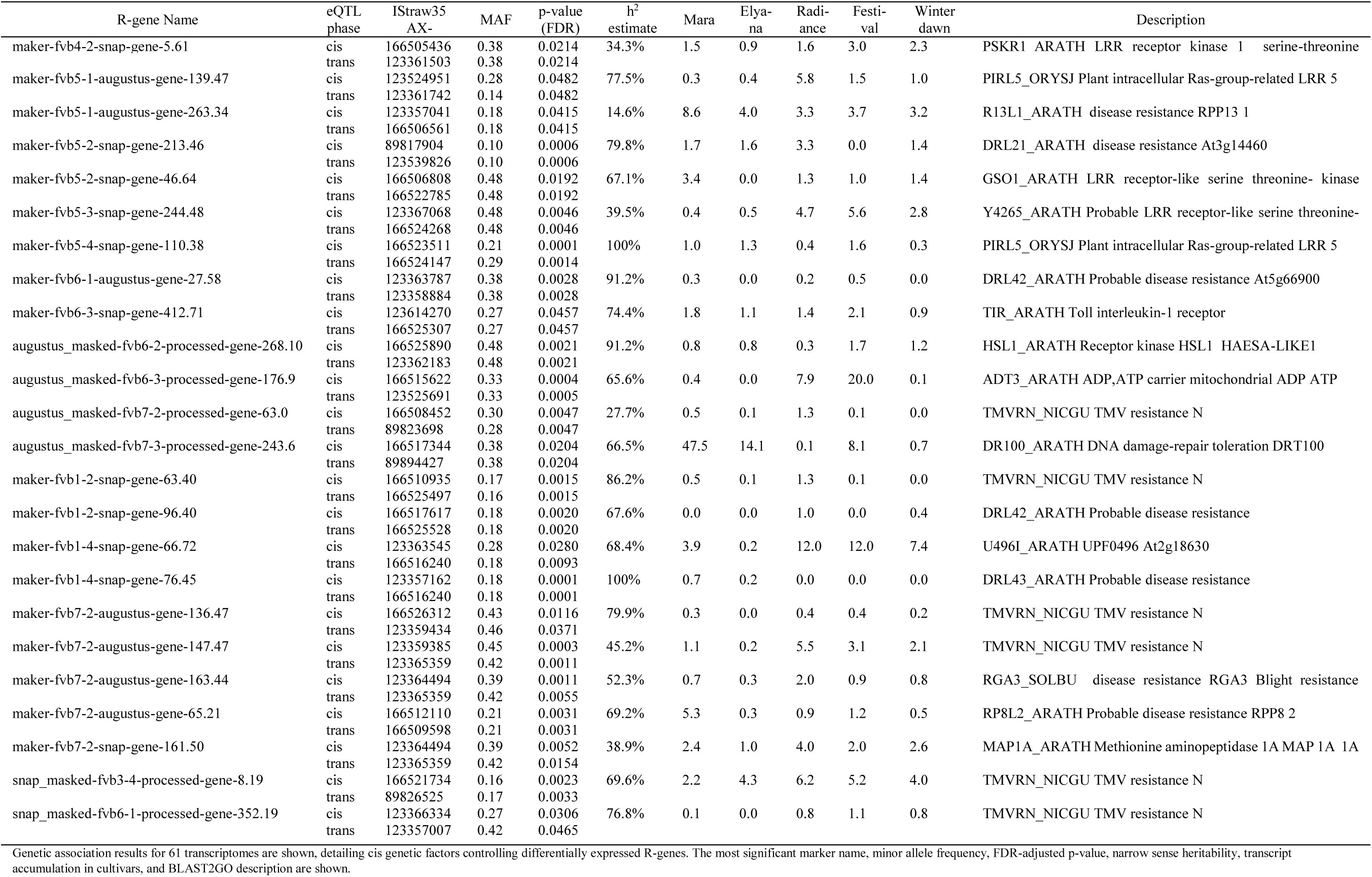
Trans and Cis Expression-QTL for Strawberry Fruit

A representative eQTL R-gene (maker-Fvb5-1-snap-gene-191.37) is detailed in Figure 6. Analysis by BLAST2GO indicates an “RPM1-like disease resistance gene” whose *Arabidopsis thaliana* homolog confers resistance to *Pseudomonas syringae*. This gene is hereafter referred to as “*FaRPM1.1*”. An unusual double NB-ARC structure is predicted for *FaRPM1.1* (Figure 6A). An eQTL was detected for this gene relative to chromosome 5 on the *F. vesca* genome position (Figure 6B). This eQTL is superficially analogous to the physical position of octoploid *FaRPM1.1* on chromosome 5, homoeolog 1 (Figure 6C). The significant eQTL markers are not included in the ‘Holiday’ × ‘Korona’ octoploid genetic map, impeding a recombination-based subgenomic genetic association (van Dijk *et al.*, 2014). However, the associated marker physical sequences match uniquely to the ‘Camarosa’ chromosome 5 homoeolog 1 locus, within several kilobases of the *FaRPM1.1*, confirming a *cis*-eQTL designation (Figure 6D).

**Figure 5.**
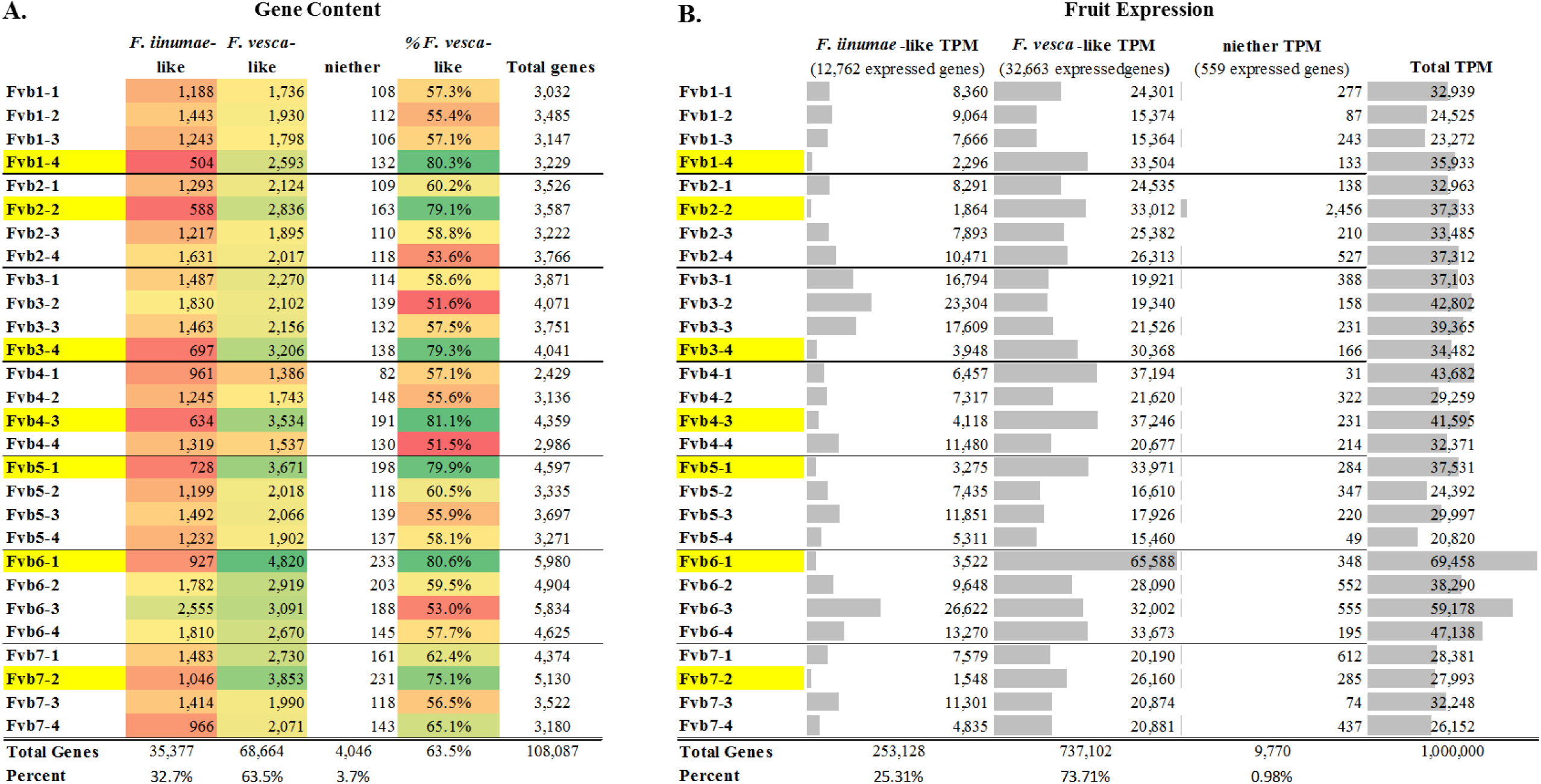
General Retention and Expression Bias in Octoploid Strawberry. ‘Camarosa’ gene models are categorized as either more *F. vesca*-like, more *F. iinumae*-like, or neither. Red-green color scale indicates low-to-high gene content, respectively. Yellow highlight indicates the most *F. vesca-*like homoeolog. **A**. Gene content per homeologous chromosome, by putative ancestral gene similarity. **B.** Relative transcript accumulation of all genes in the fruit, by putative ancestral similarity.

**Figure 6.**
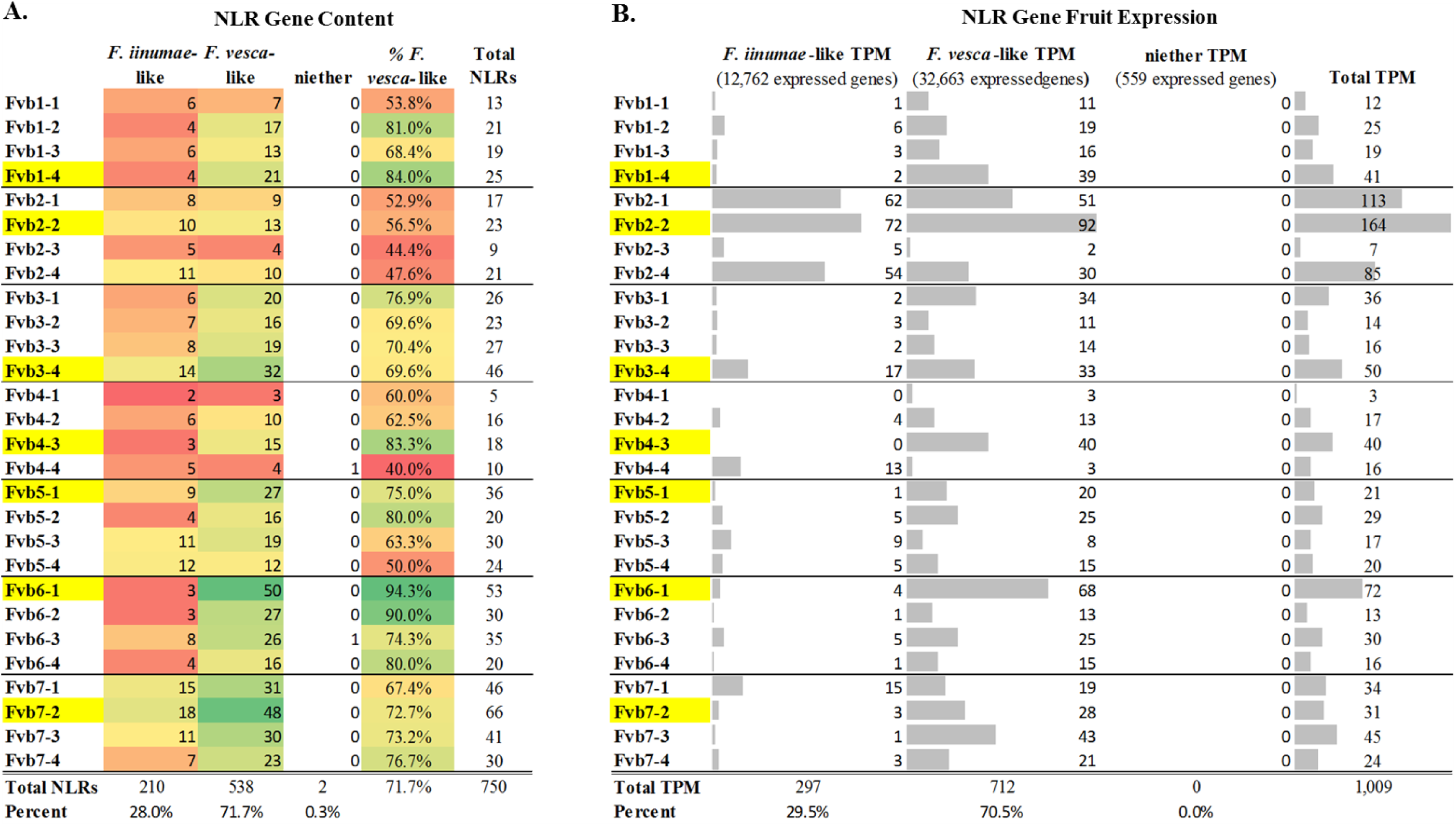
NLR-gene Retention and Expression Bias in Octoploid Strawberry. ‘Camarosa’ gene models are categorized as either more *F. vesca*-like, more *F. iinumae*-like, or neither. Red-green color scale indicates low-to-high gene content, respectively. Yellow highlight indicates the most *F. vesca*-like homoeolog. **A**. NLR-gene content per homeologous chromosome, by putative ancestral gene similarity. **B.** Relative transcript accumulation of NLR-genes in the fruit, by putative ancestral similarity.

### Evolutionary Pressure on *F. ×ananassa* R-genes

Elevated median *d*N/*d*S ratios were observed across all 1,962 predicted *F.* ***×*** *ananassa* R-genes (0.4232) compared to non R-genes (0.3526) (Figure 4). Fewer R-genes exhibited extremely low *d*N/*d*S ratios, indicating that high degrees of R-gene conservation are less common. However, a similar rate of hypervariable genes (*d*N≫*d*S) was observed between R-genes and non R-genes. A complete list of dN/*d*S ratios for each *F. ×ananassa* R-gene is provided in Supplementary Table 1.

R-gene *d*N/*d*S values were compared against transcript accumulation across various strawberry tissues and receptacle stages. R-genes with low transcript accumulation across all tissues were correlated with higher *d*N/*d*S ratios (Pearson’s r = −0.69, p < .0001) (Figure S5). In other words, R-genes with poor evidence of expression also have higher ratios of non-synonymous mutation capable of altering amino acid sequences and affecting protein function.

### Subgenome Dominance in Octoploid Strawberry

Polyploidization is associated with rapid genome remodeling events to establish a new homeostasis, including selective gene loss and methylation. While R-gene expansiveness is often considered evolutionarily favorable, genes that are stoichiometrically or dosage sensitive are more commonly retained in duplicate after polyploidization (Edger and Pires, 2009, Birchler and Veitia, 2012, Edger *et al.*, 2017a). The ‘Camarosa’ octoploid genome, in comparison with the genomes from its diploid *F. vesca*-like and *F. iinumae*-like ancestors, has provided an ideal platform to study the general biological phenomena of post-hybridization genome remodeling and subgenome dominance (Edger *et al.*, 2019). To gauge R-gene post-hybridization retention specifically, a gene-focused baseline assessment of subgenome dominance in the ‘Camarosa’ octoploid genome was necessary. Putative gene ancestry was predicted based on gene-by-gene sequence comparisons to determine the closest ‘Camarosa’ gene homologs in *F. vesca* (Fragaria_vesca_v2.0.a2.cds) and *F. iinumae* (FII_r1.1cds), which is representative of the highly similar ‘old world’ subgenomes. This gene-by-gene putative orthology analysis was selected over a total comparison of homeologous chromosomes, as extensive genetic transfer from the *F. vesca*-like subgenome has strongly converted all subgenomes to contain *F. vesca*-like genes over time (Tennessen *et al.*, 2014), and because the *F. iinumae* FII_r1.1 genome is incompletely assembled and is not amenable to whole-genome alignment. By this facile coding-sequence comparison method, a significant bias towards the retention and/or expansion of *F. vesca*-like genes is observed in the ‘Camarosa’ genome (Figure 5A), with an even stronger bias towards *F. vesca*-like fruit gene expression (Figure 5B) consistent with previous analyses (Edger *et al.*, 2019). Of 108,087 *F. ×ananassa* ‘Camarosa’ predicted gene models, 68,664 genes (63.5%) were most similar to an *F. vesca* gene model, with 35,377 (32.7%) most similar to an *F. iinumae* gene model, with a minority of genes not closely matching either. A single homoeologous chromosome with significantly more *F. vesca*-like genes (∼80% *F. vesca*-like) was seen in every chromosomal group. In a majority of cases, this putative *F. vesca*-derived chromosome possesses the greatest total gene content of the chromosome group. In 61 fruit transcriptomes, 73.7% of total transcripts derived from a gene sequence most similar to *F. vesca*, corresponding to a 10.2% expression increase relative to the baseline genomic retention bias. This bias towards the expression of *F. vesca*-like sequences was seen on every subgenome (Figure 5B, yellow highlight).

### NLR-gene Subgenome Dominance

Significant gene retention bias towards R-genes that are more *F. vesca*-like is observed in ‘Camarosa’ gene models (Figure 6). Of the 750 predicted NLR-gene models, 71.7% more closely resemble a *F. vesca* gene rather than an *F. iinumae* gene (Figure 6A). This is somewhat higher than the baseline retention bias towards *F. vesca*-like genes in octoploid (63.8%) from this analysis. In every chromosome group, the *F. vesca*-like homoeologous chromosomes (yellow highlight) retained the greatest number of NLRs. Of expressed R-genes, 1,125 demonstrate the highest sequence identity with an *F. vesca* gene, 444 show highest sequence identity with an *F. iinumae* gene, and 2 (an RPW-only gene, and an LRR_8-only gene) are without significant matches to either diploid genome. While *F. vesca*-like genes contribute the most to total mature fruit NLR expression (70.5% of transcripts), this is proportional to *F. vesca*-like NLR genome content (71.3%) and is similar in magnitude to general *F. vesca* expression bias (73.7%) (Figure 6B). In other words, *F. vesca*-like NLR genes are retained in the octoploid genome somewhat above the baseline bias, but do not experience the additional expression bias that is a generic feature of *F. vesca-*like transcripts.

### RenSeq for Strawberry Resistance Genes

A panel of sequence capture probes was designed based on putative R-gene sequences discovered in the genomes of *F. ×ananassa* ‘Camarosa’, *F. vesca* genotype Hawaii 4, *F. iinumae*, and de novo fruit transcriptomes from *F. ×ananassa* cultivars ‘Mara des Bois’ and ‘Florida Elyana’. Benchtop RenSeq capture on genomic DNA was performed on a collection of sixteen strawberry genotypes, including twelve *F. ×ananassa* advanced breeding selections, three *F. ×ananassa* disease-resistant cultivars, and a diploid *F. vesca*. As a preliminary validation of capture efficiency with this novel RenSeq probe panel, multiplexed Illumina sequencing was performed on captured R-gene genomic libraries. An average of 2.60 million reads (2 × 100bp) was obtained for each of sixteen libraries from a single lane. Reads from octoploid and diploid lines were mapped to their respective annotated genomic references. An average R-gene resequencing depth of 26x was achieved in the ‘Camarosa’ RenSeq line and 30x in the *F. vesca*, with similar coverage ranges in the other diverse octoploid accessions (Figure 8). In the ‘Camarosa’ RenSeq line, 68% of reads mapped to an annotated resistance gene, while an additional 20% of reads mapped to a non-R-gene gene model. In *F. vesca* this efficiency was lower, where 36% of reads mapped to an annotated R-gene. A FASTA of RenSeq probes is provided for use in File S2. Example probe coverage is detailed in Figure S6.

**Figure 7.**
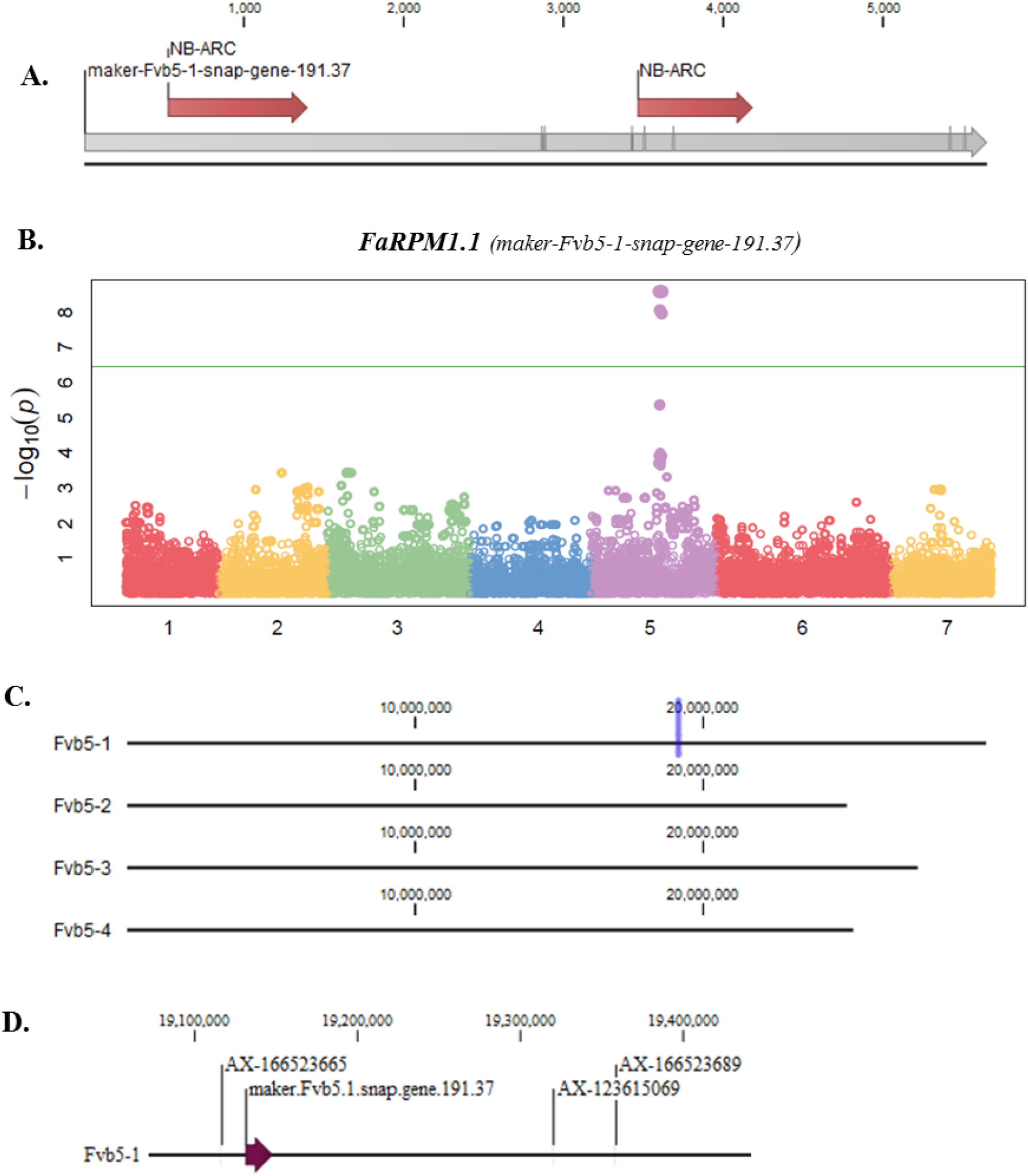
Example *cis*-eQTL of a Fruit-Expressed Strawberry R-gene. **A**. Domain analysis of the ‘Camarosa’ putative resistance gene *FaRPM1.1* indicates two NB-ARC domains. Grey lines delineate exon-exon borders in the mature predicted transcript. **B.** Octoploid fruit expression of *FaRPM1.1* associates with a single locus on chromosome 5. **C.** The *Fvb5-1* subgenomic location of *FaRPM1.1* in the octoploid ‘Camarosa’ genome is indicated (purple vertical line). **D**. Three equally highly-significant GWAS markers (p-value 8.09E-06, post-FDR adjustment) show close subgenome co-localization with *FaRPM1.1*.

**Figure 8.**
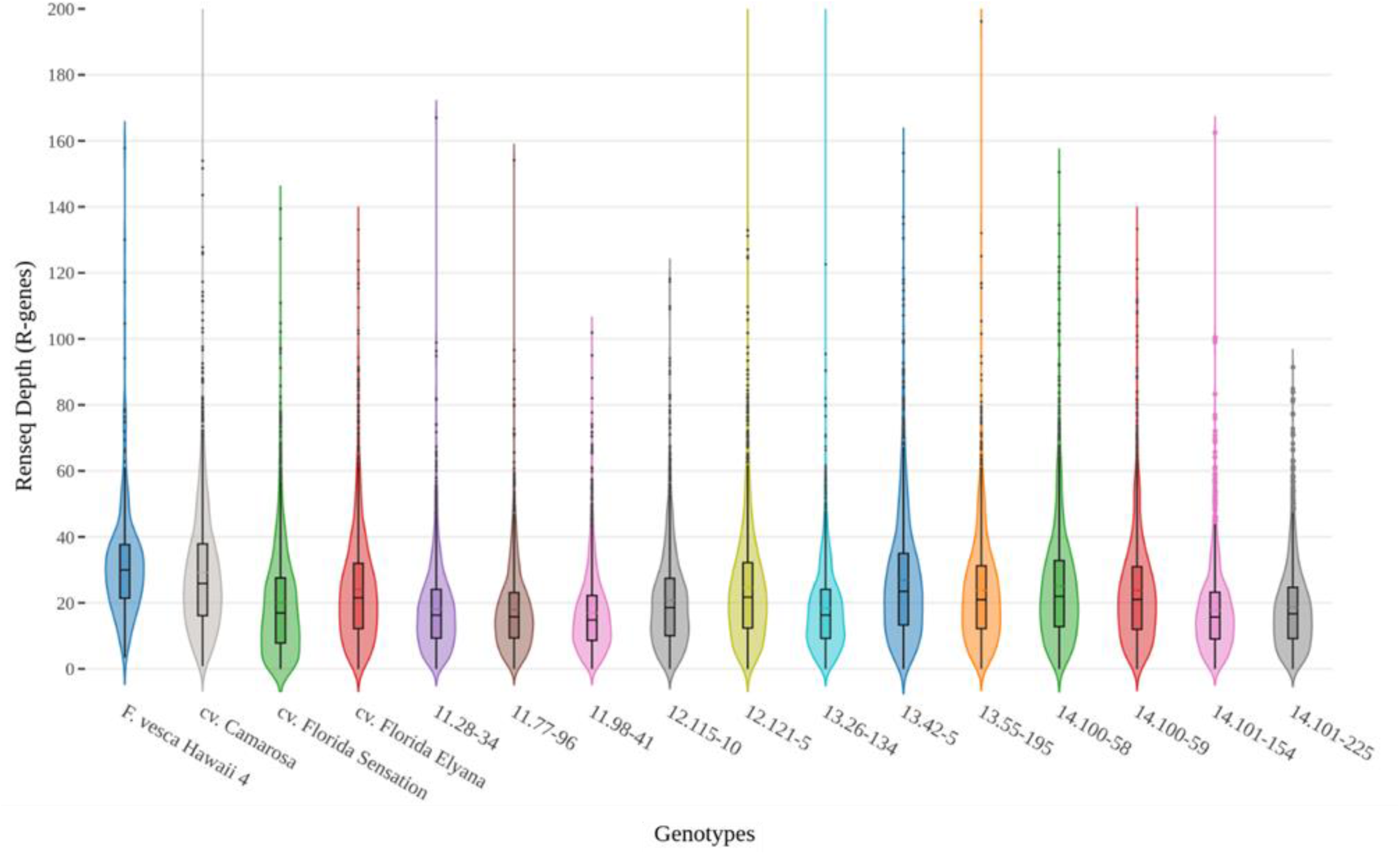
RenSeq Increases Sequencing Depth for R-gene Loci in Multiplexed Octoploid Genomes. Sixteen disease-resistant strawberry genomic libraries (fifteen octoploid accessions and diploid F. vesca) were enriched for R-genes and sequenced via Illumina Hiseq yielding an average of 2.60 million reads per genomic library. Roughly half of all sequencing reads mapped to a previously-identified R-gene locus, representing a substantial sequence enrichment relative to R-gene genomic representation.

## DISCUSSION

These results provide a characterization of the R-gene complement of cultivated octoploid strawberry and the relationship to the extant diploid relatives, *F. vesca* and *F. iinumae*. Commercial strawberry is hypothesized to contain a single *F. vesca*-like subgenome, and three highly-similar ‘old world’ subgenomes which are likely derived from *F. iinumae, F. viridis*, and *F. nipponica* (Edger *et al.*, 2019). Polyploidization is associated with massive genome remodeling events including gene loss (Edger *et al.*, 2017a, Edger *et al.*, 2018). Linkage-map comparisons in octoploid and diploid strawberry have uncovered extensive unidirectional homoeologous exchanges which have broadly converted the three *‘old world’ F. innumae*-like subgenomes to be more *F. vesca*-like (Tennessen *et al.*, 2014). This finding explains the difficulties of clear ancestral delineation of strawberry homoeologs (Vining *et al.*, 2017). Recent analysis of the octoploid genome reveals that biased homologous exchanges have converted other subgenomes to be more like the dominant *F. vesca*-like subgenome (Edger *et al.*, 2019). The present gene-level homology and expression analysis shows the majority of *F. vesca*-like dominance is derived from *F. vesca*-like genes residing on alternate subgenomes. For NLR genes in particular, the bias towards *F. vesca*-like genomic retention was more pronounced. Unlike general octoploid genes, expression of *F. vesca*-like and *F. iinumae*-like NLRs is proportional to their genomic representation. This finding provides potential insight into the practical drivers of subgenome conversion. Consolidation of redundant genes and maintenance of stoichiometrically sensitive genes has been hypothesized as a driver for gene retention bias. NLRs are involved in consequential and sensitive protein-level interactions, including signaling functions requiring homo- and hetero-dimerizations. Avoidance of dysfunctional NLR molecular interactions may have contributed to the observed biases in NLR retention and expression, post-polyploidization.

Multiple distinct NLR clades with identical domain architectures were detected, likely distinguishing intra-subgenome homologs from different subgenomes. These likely reflect broad ancestral sequence divergences prior to hybridization. Comparison of R-genes in octoploid and diploid strawberry reveals enrichment of different subtypes. The ‘Camarosa’ genome shows a large increase in complete TNL-type R-genes and a concomitant decrease in truncated TIR-only genes, relative to its diploid ancestral relatives. The *F. iinumae* genome shows a considerably larger amount of CNL-types. TNLs have been nearly eliminated from most monocot genomes in bias towards CNL-types (Nepal *et al.*, 2017). The reasons for emerging divisions in TNL/CNL content in plant genomes remains unclear. In hybrid *F. ×ananassa*, it is possible that relatively high number of complete TNL genes is a result of higher rate of retention post-polyploidization in this category. Many of the non-classical domains discovered in ‘Camarosa’ R-genes have also been found and characterized in the R-genes of other species. These include an LRR/Malectin-like RLK protein, which mediates powdery mildew resistance in barley and wheat (Rajaraman *et al.*, 2016). Atypical R-gene domains physically associated with NB-ARCs have been implicated in a variety of active disease resistance functions, including signal transduction and defense gene activation, and serving as decoy endogenous sequences to bait pathogen effectors into direct interaction and detection (Khan *et al.*, 2016).

A large proportion of strawberry NLR genes from octoploid and diploid genomes are associated with RPW8 domains. It has been suggested that the RPW8 domain emerged with the earliest land plants and subsequently merged with NLR genes, however their prevalence across plant genomes varies widely (Zhong and Cheng, 2016). The RPW domain appears to have been completely lost in monocots, and is rare in many other species. R-gene genomic studies frequently neglect to assess the presence of NLR-associated RPW domains. Two NBS-RPW8 proteins conferring mildew-resistance have been described in the *Arabidopsis thaliana* genome (Xiao *et al.*, 2001) that retained their function when expressed in grape (Hu *et al.*, 2018). The *At*RPW8.2 gene was recently shown to induce the expression of defense-related genes when expressed in strawberry leaves (Cui *et al.*, 2017). This R-gene subtype has apparently expanded in strawberry, possibly due to unusually high mildew disease pressure exerted on strawberry species and intense selection for resistance. However, R-gene domain content is not reliably predictive of resistance specificity, and close R-gene paralogs are known to confer resistance to pathogens in entirely different kingdoms (Wen *et al.*, 2015). Interestingly, the strawberry RPW8 domain is frequently found in association with NB-ARC-containing genes but never with TIRs. The purpose of RPW8 gene expansion in strawberry remains an interesting open question.

Octoploid NLR transcript accumulation is low throughout the strawberry plant, but is particularly low in the mature receptacle. This is a somewhat unexpected result due to the many pathogens targeting this susceptible organ. It is possible that only certain R genes are highly upregulated in the response to pathogen attack. Another possibility is that resistance based on the hypersensitive response may be less effective at mature stages, where cell wall disruption has already initiated with ripening and the intercellular environment is conducive to pathogen growth. Transcriptional response to *Botrytis cinerea* infection in the mature octoploid receptacle led to differential expression of over 1,500 genes, including secondary metabolism and pathogenesis-related (PR) genes, but only 15 NLR genes (Xiong *et al.*, 2018). In the present study, elevated NLR transcription in the green, white, and turning stages suggest NLR-based resistance may be more prevalent at these earlier developmental stages. The highest levels of NLR expression were seen in the roots and leaves, indicating this mode of resistance may be more common in these tissues. NLR expression overlaps poorly between tissues, supporting the concept that NLRs are optimized for each tissue. It would be interesting to examine the patterns of tissue specific expression of R-genes against different strawberry pathogens.

The genetics of differential fruit expression of R-genes in strawberry cultivars was examined via eQTL analysis. In many cases, the identified genetic markers described presence/absence of R-gene expression. The identified eQTLs were often due to a *cis* variant at a single detectable locus, very close to the physical position of the gene itself. This is suggestive of a mutation in a *cis*-regulatory element, such as the gene promoter or 5’-UTR, or a genic presence/absence structural variation. Such presence/absence variation affects nearly 20% of genes in the *Brassica oleracea* pangenome and is a major contributor of agronomic trait diversity (Golicz *et al.*, 2016). As these strawberry R-gene eQTL are derived from crosses of cultivars with differing ranges of pathogen susceptibility, these eQTL genes represent strong candidates for functional disease resistance and potential genetic improvement. These disclosed R-gene eQTL marker sequences may be cross-referenced with existing disease-resistance QTL to potentially identify causal R-genes. As categories of R-genes are expressed at very low levels unless induced by pathogens (Lai and Eulgem, 2017), the genotype × pathogen interaction may have lowered confidence values or introduced possible type II errors in eQTL detection. However, the reproducibility of *cis*-eQTL tends to be particularly high in related populations (Peirce *et al.*, 2006). Additional replicates and infected/non-infected challenge conditions will likely reveal additional eQTL associations and greatly improve the confidence of heritability estimates, and may be used to validate pathogen-induced R-gene candidates.

*F. ×ananassa* predicted R-genes (NLRs and other R-gene types) have elevated average *d*N/*d*S ratios compared to non R-genes, indicating greater overall tendency towards divergent selection. R-genes with very low *d*N/*d*S ratios are likely to be conserved disease resistance genes. This active evolutionary selection is highly indicative of function. Of particular interest are strawberry R-genes demonstrating both low *d*N/*d*S values and low transcript levels across all tissues (Table S1). Many functional R-genes are expressed at low levels, either constitutively or until elicited by the proper pathogen (Lai and Eulgem, 2017). Such R-genes may be difficult to distinguish from pseudogenes on a purely transcriptional bases. Low *d*N/*d*S values demonstrate selective pressure to maintain these sequences, offering evidence of maintained function despite low expression. The results of this combinatorial analysis can be used help identify novel sources of R-gene-based resistance which may be otherwise difficult to detect. It should be noted that this analysis is performed in the context of a single cultivar, which has undergone several centuries of artificial selection. It is possible that wild octoploid species may reveal different and more natural patterns of disease-resistance selection. More sequenced accessions from geographically diverse wild and cultivated germplasm are needed. Further analysis on the octoploid pangenome will reveal more detailed selection patterns, and more importantly, reveal recent selection sweep events which may have occurred in certain R-gene groups.

Many R-genes were discovered clustered in the genomes of both octoploid and diploid strawberry, highlighting the challenges of resolving individual R-genes via association mapping and positional cloning. The difficulty of isolating functional R-genes from strawberry disease resistance QTL was the principle motivator of this analysis. A thorough identification of R-genes in the octoploid genome is necessary for future genomics and genetics analysis in strawberry disease-resistance breeding programs. Additionally, this information is prerequisite for creating a RenSeq probe panel, to facilitate targeted R-gene sequencing in breeding programs.

A novel strawberry RenSeq capture-probe library was developed based on the R-gene sequences identified from genomic and transcriptomic resources. This 39,501-probe panel was experimentally validated using octoploid and diploid genomes and resulted in an average ∼20× R-gene resequencing depth per genomic library, using only multiplexed short reads. RenSeq assembly in ‘Camarosa’ and the *F. vesca* genotype Hawaii 4 resulted in significant coverage of R-genes. Interestingly, the capture efficiency (R-gene reads over total reads) was somewhat lower in *F. vesca*, likely representing the saturation of capture probes in the smaller *F. vesca* genome. Similar rates of perfect sequence matching along the entire read in ‘Camarosa’ and *F. vesca* (66.24% and 69.68%, respectively) indicates that theoretical octoploid reference sequence errors are not likely promoting RenSeq assembly error in ‘Camarosa’. However, 14.4% of mapped ‘Camarosa’ R-gene reads have an equally valid alternative R-gene mapping locus, compared with just 4.83% in *F. vesca*. This difference indicates that homoeologous sequence redundancy is an appreciable issue for mapping short-reads in polyploids, even with an isogenic (but not haplotype-specific) mapping reference. Longer sequencing read-lengths, spanning less well-conserved non-coding sequences, will assist in de novo resolution of similar loci in octoploid strawberry. Combining RenSeq with longer-read third generation sequencing technologies will allow for improved de novo assembly of R-gene loci, and will greatly facilitate causal mutation detection within disease resistance QTL in octoploid strawberry.

## MATERIALS AND METHODS

### Plant Populations and Genetic Materials

Three pedigree-connected and segregating strawberry populations were created from crosses ‘Florida Elyana’ × ‘Mara de Bois’, ‘Florida Radiance’ × ‘Mara des Bois’, and ‘Strawberry Festival’ × ‘Winter Dawn’ (Figure S7). These cultivars and 54 progeny were selected for RNAseq and Istraw35 SNP genotyping analysis (Verma *et al.*, 2017), and were used to identify expressed genes and R-gene eQTL. De novo assemblies of ‘Mara des Bois’ and ‘Florida Elyana’ were also used to help design RenSeq capture probes.

For RenSeq, 14 disease resistant octoploid cultivars and elite breeding lines were selected from the University of Florida breeding program, and supplemented with ‘Camarosa’ and with the ancestral diploid *F. vesca*. The RenSeq lines are *F. vesca* genotype Hawaii 4, ‘Camarosa’, Sweet Sensation® ‘Florida127’, ‘Florida Elyana’, 11.28-34, 11.77-96, 11.98-41, 12.115-10, 12.121-5, 13.26-134, 13.42-5, 13.55-195, 14.100-58, 14.100-59, 14.101-154, and 14.101-225.

### Identification of R-genes in Strawberry spp.

R-genes were predicted from the strawberry octoploid ‘Camarosa’ draft genome “F_ana_Camarosa_6-28-17.rm” (Edger *et al.*, 2019), the diploid *F. vesca* reassembly “Fragaria_vesca_v2.0.a2” (Tennessen *et al.*, 2014), and the diploid *F. innumae* assembly “FII_r1.1” (Hirakawa *et al.*, 2014). Domain-level analysis was performed using the CLC Genomics Workbench 11 HMM implementation to search for Pfam-v29 domains on translated gene models from all genomic and transcriptomic strawberry resources. Motif search was performed on all translated gene models, using 56 R-gene-associated motifs collected from (Lukasik and Takken, 2009, Jupe *et al.*, 2012, Van Ghelder and Esmenjaud, 2016). The CLC Genomics Workbench 11 (CLC Bio, Denmark) pattern discovery tool was trained on a preliminary list of strawberry R-genes, and novel motifs were reiterated back to all protein models. The Ncoils sequence analysis algorithm (Lupas *et al.*, 1991) was used to detect coiled-coil domains, and the output was parsed into GFF3 format for protein list reannotation. BLAST2GO annotation (Conesa *et al.*, 2005) was performed to assign putative functions to all genes and confirm sequence association with disease resistance in a cross-referenced database.

Protein models containing canonical R-gene domains (eg. NB-ARC domain) were selected for inclusion as R-genes, as were gene models with more common domains (eg. LRR) with supporting evidence of an R-gene-associated motif. BLAST2GO annotated disease resistance associated genes not meeting the domain and motif-level criteria were analyzed for potential inclusion, leading to the inclusion of many LRR-containing RLK putative R-genes.

### NB-ARC Phylogenetic Analysis

NB-ARC domains were extracted from *F. iinumae, F. vesca*, and *F. ×ananassa* ‘Camarosa’. The CIPRES Science Gateway (Miller *et al.*, 2010) was utilized for full-length protein sequence alignment using MUSCLE v3.7 (Edgar, 2004) and Maximum likelihood analysis using RAxML v8.2.10 (Stamatakis, 2014). Tree construction was performed using the PROTGAMMA rate distribution model with 100 bootstrap replicates, and rooted with human APAF-1. This process was replicated five times using different random number seeds. Trees were visualized in CLC Genomic Workbench 11 with a 50% threshold bootstrap value. Word clouds were generated per clade based on the relative domain content of the full proteins.

### Subgenome Dominance in Octoploid Strawberry R-genes

The closest homolog for each *F. ×ananassa* ‘Camarosa’ gene in either Fragaria_vesca_v2.0.a2.cds or FII_r1.1cds was determined via BLAST analysis (e-value threshold < 0.1, word size=25, match=1, mismatch=l, existence=0, extension=2). *F. vesca*-like and *F. iinumae*-like gene counts and TPMs were independently calculated for each octoploid chromosome. This process was performed first on all genes in the ‘Camarosa’ genome to establish the baseline gene retention and expression bias, and then repeated using only predicted NLR genes.

### *d*N/*d*S Analysis

*d*N and *d*S values were computed using a set of custom scripts (https://github.com/Aeyocca/ka_ks_pipe/). Orthologous genes between the *F. ×ananassa and F. vesca* v4 (Edger *et al.*, 2017b) genomes were identified using the compara module in JCVI utilities library (Tang et al., 2015). Filtering of the JCVI utilities output was performed using a custom perl script to identify the best syntenic ortholog and best blast hit below e-value 1e-4. Alignment of each orthologous gene pair was performed using MUSCLE v3.8.31 (Edgar, 2004), followed by PAL2NAL (v14) (Suyama *et al.*, 2006) to convert the peptide alignment to a nucleotide alignment. Finally, *d*N and *d*S values were computed between those gene pairs using codeml from PAML Version 4.9h (Yang, 2007) with parameters specified in the control file found in the GitHub repository listed above.

### Tissue-specific Transcriptome Analysis

Raw short read RNAseq libraries from various ‘Camarosa’ tissue (Sánchez-Sevilla *et al.*, 2017) with the study reference PRJEB12420 were download from the European Nucleotide Archive (https://www.ebi.ac.uk/ena). The complete 54 library RNAseq experiment consisted of six independent green receptacle libraries, six white receptacle libraries, six turning receptacle libraries, six red receptacle libraries, three root libraries, three leaf libraries, and six achene libraries each for all corresponding fruit stages. Raw RNAseq reads were assembled to the ‘Camarosa’ reference using the same pipeline as previously described for fruit transcriptome population analysis. Expression values from biologically-replicated libraries were averaged. Clustvis (Metsalu and Vilo, 2015) was used for tissue-based RNAseq clustering and heatmap visualization using correlation distance and average linkage with scaling applied using default parameters.

### Fruit Transcriptome Analysis

61 fruit transcriptomes were sequenced via Illumina paired-end RNAseq (Avg. 65million reads, 2×100bp), and consisted of parents and progeny from crosses of ‘Florida Elyana’ × ‘Mara de Bois’, ‘Florida Radiance’ × ‘Mara des Bois’, and ‘Strawberry Festival’ × ‘Winter Dawn’. Reads were trimmed and mapped to the *F. ×ananassa* octoploid ‘Camarosa’ annotated genome using CLC Genomic Workbench 11 (mismatch cost of 2, insertion cost of 3, deletion cost of 3, length fraction of 0.8, similarity fraction of 0.8, 1 maximum hit per read). Reads that mapped equally well to more than one locus were discarded from the analysis. RNAseq counts were calculated in Transcripts Per Million (TPM). Three-dimensional principle component analysis (PCA) was performed on all RNAseq assemblies, including two replicates of ‘Mara des Bois’ fruit harvested three years apart and sequenced independently (Figure S2). Transcript abundances were normalized via the Box-Cox transformation algorithm performed in R (R. Development Core Team, 2014) prior to eQTL analysis. The BLAST2GO pipeline was used to annotate the full ‘Camarosa’ predicted gene complement.

### Genotyping and Genetic Association of Octoploid Fruit R-genes

The Affymetrix IStraw35 Axiom^®^ SNP array (Verma *et al.*, 2017) was used to genotype 60 individuals, including six parental lines from three independent biparental RNAseq populations (Figure S7). Sequence variants belonging to the Poly High Resolution (PHR) and No Minor Homozygote (NMH) marker classes were included for association mapping. Mono High Resolution (MHR), Off-Target Variant (OTV), Call Rate Below Threshold (CRBT), and Other marker quality classes, were discarded and not used for mapping. Individual marker calls inconsistent with Mendelian inheritance from parental lines were removed. The *F. vesca* physical map was used to orient marker positions as current octoploid maps do not include a majority of the available IStraw35 markers. A genome-wide analysis study (GWAS) was performed using GAPIT v2 (Tang *et al.*, 2016) performed in R. R-gene eQTL were evaluated for significance based on the presence of multiple co-locating markers of p-value < 0.05 after false discovery rate correction for multiple comparisons. *Cis* vs *trans* eQTL determinations were made by corroborating known ‘Camarosa’ physical gene position with the eQTL position the *F. vesca* map. In the example case of *FaRPM1.1*, subgenomic localization was confirmed via BLAST of the associated markers to the correct ‘Camarosa’ homoeologous chromosome.

### RenSeq Probe Design and Validation

A panel of 39,501 of 120mer-length capture probes were designed based on the set of discovered strawberry R-genes from *F. ×ananassa* ‘Camarosa’, *F. vesca* genotype Hawaii 4, *F. iinumae* genomes, and de novo fruit transcriptomes from *F. ×ananassa* ‘Mara des Bois’ and ‘Florida Elyana’. A proprietary algorithm was used to select for capture probes of ideal hybridization thermodynamics and screened for potential off-target capture in the intergenic regions of ‘Camarosa’ and *F. vesca* (Rapid Genomics LLC, Gainesville FL). Probes were designed to not span exon-exon junction, to facilitate cross-utility for both genomic and cDNA libraries (Figure S6). A minimum baseline of 1x probe coverage was provided across the length of every predicted R-gene coding sequence, and additional probes were designed against conserved R-gene domains in order to promote capture of unknown and divergent R-genes across diverse octoploid accessions. RenSeq capture was performed on genomic libraries from fifteen octoploid disease-resistant cultivars and advanced breeding selections, and *F. vesca*, based on conditions set by (Jupe *et al.*, 2014), with optimizations provided by Rapid Genomics LLC. Captured libraries were sequenced via 16× multiplexed Illumina Hiseq (2 × 100bp) and mapped to their respective annotated genomic references using CLC Genomic Workbench 11 (CLCBio, Aarhus, Denmark) (Similarity fraction = 0.9, Length fraction = 0.9, Match score = 1, Mismatch cost = 2, Insertion cost = 3, Deletion cost = 3).

### Data Availability

Supplementary figures, tables, files, and raw data are available at FigShare. Custom scripts used for performing *d*N/*d*S analysis are available at Github: https://github.com/Aeyocca/ka_ks_pipe/. Raw short read RNAseq data from fruit transcriptomes are available from the NCBI Short Read Archive under project SRP039356 (http://www.ncbi.nlm.nih.gov/sra/?term=SRP039356). Raw short read RNAseq data from the ‘Camarosa’ gene expression atlas (Sánchez-Sevilla et al., 2017) are available at the European Nucleotide Archive (https://www.ebi.ac.uk/ena) with the study reference PRJEB12420. Results derived from these data are compiled in Table S1. File S1 contains BED files for annotating the octoploid genome (Edger *et al.*, 2019) with R-genes and R-gene domains. Renseq probe sequences are provided in File S2. Istraw35 markers, map positions, and sample genotypes used in eQTL analysis are available in File S3.

## ACKNOWLEDGEMENTS

We acknowledge Aristotole Koukoulidis for ncoils prediction and annotation, Matthew Robinson for assistance with data normalization, Rapid Genomics LLC for technical assistance in probe generation and sequence capture, Anne Schwartz, Max Hogshead, Kiran Sharma and Nadia Mourad for assistance in data compilation, Ben Harrison and Max Hogshead for assistance in gDNA isolation of eQTL lines, and Dr. Alan Chambers and Dr. Jeremy Pillet for RNA isolation and RNAseq line selection. This research is supported by grants to SJK, VMW, and SL from the United Stated Department of Agriculture (http://dx.doi.org/10.13039/100000199) National Institute of Food and Agriculture (NIFA) Specialty Crops Research Initiative (#2017-51181-26833) and to SJK from the California Strawberry Commission (http://dx.doi.org/10.13039/100006760).

## LITERATURE CITED

Amil-Ruiz, F., Departamento de Bioquímica y Biología Molecular e Instituto Andaluz de Biotecnología, C.d.E.I.A.C.C.d.R.E.S.O.U.d.C.C.S., Blanco-Portales, R., Muñoz-Blanco, J. and Caballero, J.L. (2018) The Strawberry Plant Defense Mechanism: A Molecular Review. Plant and Cell Physiology, 52, 1873–1903 10.1093/pcp/pcr136.

Anciro, A., Mangandi, J., Verma, S., Peres, N., Whitaker, V.M. and Lee, S. (2018) FaRCg1: a quantitative trait locus conferring resistance to Colletotrichum crown rot caused by Colletotrichum gloeosporioides in octoploid strawberry. Theoretical and Applied Genetics 10.1007/s00122-018-3145-z.

Andolfo, G., Jupe, F., Witek, K., Etherington, G.J., Ercolano, M.R. and Jones, J.D.G. (2014) Defining the full tomato NB-LRR resistance gene repertoire using genomic and cDNA RenSeq. BMC Plant Biology, 14, 120 10.1186/1471-2229-14-120.

Arya, P., Kumar, G., Acharya, V. and Singh, A.K. (2014) Genome-Wide Identification and Expression Analysis of NBS-Encoding Genes in Malus x domestica and Expansion of NBS Genes Family in Rosaceae. PLOS ONE, 9, e107987. 10.1371/journal.pone.0107987.

Baumgartner, I.O., Patocchi, A., Frey, J.E., Peil, A. and Kellerhals, M. (2015) Breeding Elite Lines of Apple Carrying Pyramided Homozygous Resistance Genes Against Apple Scab and Resistance Against Powdery Mildew and Fire Blight. Plant Molecular Biology Reporter, 33, 1573–1583 10.1007/s11105-015-0858-x.

Birchler, J.A. and Veitia, R.A. (2012) Gene balance hypothesis: Connecting issues of dosage sensitivity across biological disciplines. Proceedings of the National Academy of Sciences, 109, 14746 10.1073/pnas.1207726109.

Conesa, A., Götz, S., García-Gómez, J.M., Terol, J., Talón, M. and Robles, M. (2005) Blast2GO: a universal tool for annotation, visualization and analysis in functional genomics research. Bioinformatics, 21, 3674–3676 10.1093/bioinformatics/bti610.

Cordova, L.G., Amiri, A. and Peres, N.A. (2017) Effectiveness of fungicide treatments following the Strawberry Advisory System for control of Botrytis fruit rot in Florida. Crop Protection, 100, 163–167 https://doi.org/10.1016/j.cropro.2017.07.002.

Cui, M.-Y., Wei, W., Gao, K., Xie, Y.-G., Guo, Y. and Feng, J.-Y. (2017) A rapid and efficient Agrobacterium-mediated transient gene expression system for strawberry leaves and the study of disease resistance proteins. Plant Cell, Tissue and Organ Culture (PCTOC), 131, 233–246 10.1007/s11240-017-1279-3.

Djian-Caporalino, C., Palloix, A., Fazari, A., Marteu, N., Barbary, A., Abad, P., Sage-Palloix, A.-M., Mateille, T., Risso, S., Lanza, R., Taussig, C. and Castagnone-Sereno, P. (2014) Pyramiding, alternating or mixing: comparative performances of deployment strategies of nematode resistance genes to promote plant resistance efficiency and durability. BMC Plant Biology, 14, 53 10.1186/1471-2229-14-53.

Edgar, R.C. (2004) MUSCLE: multiple sequence alignment with high accuracy and high throughput. Nucleic acids research, 32, 1792–1797

Edger, P.P., McKain, M.R., Bird, K.A. and VanBuren, R. (2018) Subgenome assignment in allopolyploids: challenges and future directions. Current Opinion in Plant Biology, 42, 76–80 https://doi.org/10.1016/j.pbi.2018.03.006.

Edger, P.P. and Pires, J.C. (2009) Gene and genome duplications: the impact of dosage-sensitivity on the fate of nuclear genes. Chromosome Research, 17, 699 10.1007/s10577-009-9055-9.

Edger, P.P., Poorten, T.J., VanBuren, R., Hardigan, M.A., Colle, M., McKain, M.R., Smith, R.D., Teresi, S.J., Nelson, A.D.L., Wai, C.M., Alger, E.I., Bird, K.A., Yocca, A.E., Pumplin, N., Ou, S., Ben-Zvi, G., Brodt, A., Baruch, K., Swale, T., Shiue, L., Acharya, C.B., Cole, G.S., Mower, J.P., Childs, K.L., Jiang, N., Lyons, E., Freeling, M., Puzey, J.R. and Knapp, S.J. (2019) Origin and evolution of the octoploid strawberry genome. Nature Genetics, 51, 541–547 10.1038/s41588-019-0356-4.

Edger, P.P., Smith, R., McKain, M.R., Cooley, A.M., Vallejo-Marin, M., Yuan, Y., Bewick, A.J., Ji, L., Platts, A.E., Bowman, M.J., Childs, K.L., Washburn, J.D., Schmitz, R.J., Smith, G.D., Pires, J.C. and Puzey, J.R. (2017a) Subgenome Dominance in an Interspecific Hybrid, Synthetic Allopolyploid, and a 140-Year-Old Naturally Established Neo-Allopolyploid Monkeyflower. The Plant Cell, 29, 2150

Edger, P.P., VanBuren, R., Colle, M., Poorten, T.J., Wai, C.M., Niederhuth, C.E., Alger, E.I., Ou, S., Acharya, C.B., Wang, J., Callow, P., McKain, M.R., Shi, J., Collier, C., Xiong, Z., Mower, J.P., Slovin, J.P., Hytönen, T., Jiang, N., Childs, K.L. and Knapp, S.J. (2017b) Single-molecule sequencing and optical mapping yields an improved genome of woodland strawberry (Fragaria vesca) with chromosome-scale contiguity. GigaScience, 7 10.1093/gigascience/gix12.

Farzaneh, M., Kiani, H., Sharifi, R., Reisi, M. and Hadian, J. (2015) Chemical composition and antifungal effects of three species of Satureja (S. hortensis, S. spicigera, and S. khuzistanica) essential oils on the main pathogens of strawberry fruit. Postharvest Biology and Technology, 109, 145–151 https://doi.org/10.1016/j.postharvbio.2015.06.014.

Funk, A., Galewski, P. and McGrath, J.M. (2018) Nucleotide-binding resistance gene signatures in sugar beet, insights from a new reference genome. The Plant Journal, 95, 659–671 10.1111/tpj.13977.

Golicz, A.A., Bayer, P.E., Barker, G.C., Edger, P.P., Kim, H., Martinez, P.A., Chan, C.K.K., Severn-Ellis, A., McCombie, W.R., Parkin, I.A.P., Paterson, A.H., Pires, J.C., Sharpe, A.G., Tang, H., Teakle, G.R., Town, C.D., Batley, J. and Edwards, D. (2016) The pangenome of an agronomically important crop plant Brassica oleracea. Nature Communications, 7, 13390 10.1038/ncomms13390 https://www.nature.com/articles/ncomms13390#supplementary-information.

Hammond-Kosack, K.E. and Jones, J.D.G. (1997) Plant Disease Resistance Genes. Annual Review of Plant Physiology and Plant Molecular Biology, 48, 575–607 10.1146/annurev.arplant.48.1.575.

Herrington, M.E., Hardner, C., Wegener, M., Woolcock, L.L. and Dieters, M.J. (2011) Rain Damage to Strawberries Grown in Southeast Queensland: Evaluation and Genetic Control. HortScience, 46, 832–837

Hirakawa, H., Shirasawa, K., Kosugi, S., Tashiro, K., Nakayama, S., Yamada, M., Kohara, M., Watanabe, A., Kishida, Y., Fujishiro, T., Tsuruoka, H., Minami, C., Sasamoto, S., Kato, M., Nanri, K., Komaki, A., Yanagi, T., Guoxin, Q., Maeda, F., Ishikawa, M., Kuhara, S., Sato, S., Tabata, S. and Isobe, S.N. (2014) Dissection of the Octoploid Strawberry Genome by Deep Sequencing of the Genomes of Fragaria Species. DNA Research, 21, 169–181 10.1093/dnares/dst049.

Hu, Y., Li, Y., Hou, F., Wan, D., Cheng, Y., Han, Y., Gao, Y., Liu, J., Guo, Y., Xiao, S., Wang, Y. and Wen, Y.-Q. (2018) Ectopic expression of Arabidopsis broad-spectrum resistance gene RPW8.2 improves the resistance to powdery mildew in grapevine (Vitis vinifera). Plant Science, 267, 20–31 https://doi.org/10.1016/j.plantsci.2017.11.005.

Jia, Y., Yuan, Y., Zhang, Y., Yang, S. and Zhang, X. (2015) Extreme expansion of NBS-encoding genes in Rosaceae. BMC Genetics, 16,48 10.1186/s12863-015-0208-x.

Jupe, F., Chen, X., Verweij, W., Witek, K., Jones, J.D.G. and Hein, I. (2014) Genomic DNA Library Preparation for Resistance Gene Enrichment and Sequencing (RenSeq) in Plants. In Plant-Pathogen Interactions: Methods and Protocols (Birch, P., Jones, J.T. and Bos, J.I.B. eds).Totowa, NJ: Humana Press, pp. 291–303.

Jupe, F., Pritchard, L., Etherington, G.J., MacKenzie, K., Cock, P.J.A., Wright, F., Sharma, S.K., Bolser, D., Bryan, G.J., Jones, J.D.G. and Hein, I. (2012) Identification and localisation of the NB-LRR gene family within the potato genome. BMC Genomics, 13, 75 10.1186/1471-2164-13-75.

Khan, M., Subramaniam, R. and Desveaux, D. (2016) Of guards, decoys, baits and traps: pathogen perception in plants by type III effector sensors - ScienceDirect. Current Opinion in Microbiology, 29, 49–55 10.1016/j.mib.2015.10.006.

Lai, Y. and Eulgem, T. (2017) Transcript-level expression control of plant NLR genes. Molecular Plant Pathology, 19, 1267–1281 10.1111/mpp.12607.

Lukasik, E. and Takken, F.L.W. (2009) STANDing strong, resistance proteins instigators of plant defence. Current opinion in plant biology, 12, 427–436

Lupas, A., Van Dyke, M. and Stock, J. (1991) Predicting Coiled Coils from Protein Sequences. Science, 252, 1162–1164

Mangandi, J., Verma, S., Osorio, L., Peres, N.A., van de Weg, E. and Whitaker, V.M. (2017) Pedigree-Based Analysis in a Multiparental Population of Octoploid Strawberry Reveals QTL Alleles Conferring Resistance to *Phytophthora cactorum*. G3: Genes|Genomes|Genetics, 7, 1707

Metsalu, T. and Vilo, J. (2015) ClustVis: a web tool for visualizing clustering of multivariate data using Principal Component Analysis and heatmap. Nucleic Acids Research, 43, W566–W570 10.1093/nar/gkv468.

Miller, M.A., Pfeiffer, W. and Schwartz, T. (2010) Creating the CIPRES Science Gateway for inference of large phylogenetic trees. In 2010 Gateway Computing Environments Workshop (GCE), pp. 1–8.

Nepal, M.P.A. E. J., Neupane, S. and Benson, B.V. (2017) Comparative Genomics of Non-TNL Disease Resistance Genes from Six Plant Species. In Genes (Basel).

Peirce, J.L., Li, H., Wang, J., Manly, K.F., Hitzemann, R.J., Belknap, J.K., Rosen, G.D., Goodwin, S., Sutter, T.R., Williams, R.W. and Lu, L. (2006) How replicable are mRNA expression QTL? Mammalian Genome, 17, 643–656 10.1007/s00335-005-0187-8.

Pincot, D.D.A., Poorten, T.J., Hardigan, M.A., Harshman, J.M., Acharya, C.B., Cole, G.S., Gordon, T.R., Stueven, M., Edger, P.P. and Knapp, S.J. (2018) Genome-Wide Association Mapping Uncovers *Fw1*, a Dominant Gene Conferring Resistance to Fusarium Wilt in Strawberry. G3: Genes|Genomes|Genetics, 8, 1817 10.1534/g3.118.200129.

R. Development Core Team (2014) R: A language and environment for statistical computing.

Rajaraman, J., Douchkov, D., Hensel, G., Stefanato, F.L., Gordon, A., Ereful, N., Caldararu, O.F., Petrescu, A.J., Kumlehn, J., Boyd, L.A. and Schweizer, P. (2016) An LRR/Malectin Receptor-Like Kinase Mediates Resistance to Non-adapted and Adapted Powdery Mildew Fungi in Barley and Wheat. Front Plant Sci, 7 10.3389/fpls.2016.01836.

Roach, J.A., Verma, S., Peres, N.A., Jamieson, A.R., van de Weg, W.E., Bink, M.C.A.M., Bassil, N.V., Lee, S. and Whitaker, V.M. (2016) FaRXf1: a locus conferring resistance to angular leaf spot caused by Xanthomonas fragariae in octoploid strawberry. Theoretical and Applied Genetics, 129, 1191–1201 10.1007/s00122-016-2695-1.

Salinas, N., Verma, S., Peres, N. and Whitaker, V.M. (2018) FaRCa1: a major subgenome-specific locus conferring resistance to Colletotrichum acutatum in strawberry. Theoretical and Applied Genetics 10.1007/s00122-018-263-7.

Stamatakis, A. (2014) RAxML version 8: a tool for phylogenetic analysis and post-analysis of large phylogenies. Bioinformatics, 30, 1312–1313 10.1093/bioinformatics/btu033.

Suyama, M., Torrents, D. and Bork, P. (2006) PAL2NAL: robust conversion of protein sequence alignments into the corresponding codon alignments. Nucleic acids research, 34, W609–W612

Sánchez-Sevilla, J.F., Vallarino, J.G., Osorio, S., Bombarely, A., Posé, D., Merchante, C., Botella, M.A., Amaya, I. and Valpuesta, V. (2017) Gene expression atlas of fruit ripening and transcriptome assembly from RNA-seq data in octoploid strawberry (Fragaria×ananassa). Scientific Reports, 7, 13737 10.1038/s41598-017-14239-6.

Tang, Y., Liu, X., Wang, J., Li, M., Wang, Q., Tian, F., Su, Z., Pan, Y., Liu, D., Lipka, A.E., Buckler, E.S. and Zhang, Z. (2016) GAPIT Version 2: An Enhanced Integrated Tool for Genomic Association and Prediction. The Plant Genome, 9 10.3835/plantgenome2015.11.0120.

Tennessen, J.A., Govindarajulu, R., Ashman, T.-L. and Liston, A. (2014) Evolutionary Origins and Dynamics of Octoploid Strawberry Subgenomes Revealed by Dense Targeted Capture Linkage Maps. Genome Biology and Evolution, 6, 3295–3313 10.1093/gbe/evu261.

van Dijk, T., Pagliarani, G., Pikunova, A., Noordijk, Y., Yilmaz-Temel, H., Meulenbroek, B., Visser, R.G.F. and van de Weg, E. (2014) Genomic rearrangements and signatures of breeding in the allo-octoploid strawberry as revealed through an allele dose based SSR linkage map. BMC Plant Biology, 14, 55 10.1186/1471-2229-14-55.

Van Ghelder, C. and Esmenjaud, D. (2016) TNL genes in peach: insights into the post-LRR domain. BMC Genomics, 17, 317 10.1186/s12864-016-2635-0.

Verma, S., Bassil, N.V., van de Weg, E., Harrison, R.J., Monfort, A., Hidalgo, J.M., Amaya, I., Denoyes, B., Mahoney, L., Davis, T.M., Fan, Z., Knapp, S. and Whitaker, V.M. (2017) Development and evaluation of the Axiom® IStraw35 384HT array for the allo-octoploid cultivated strawberry Fragaria ×ananassa: International Society for Horticultural Science (ISHS), Leuven, Belgium, pp. 75–82.

Verma, S., Osorio, L.F., Lee, S., Bassil, N.V. and Whitaker, V.M. (2018) Genome-Assisted Breeding in the Octoploid Strawberry. In The Genomes of Rosaceous Berries and Their Wild Relatives (Hytönen, T., Graham, J. and Harrison, R. eds). Cham: Springer International Publishing, pp. 161–184.

Vining, K.J., Salinas, N., Tennessen, J.A., Zurn, J.D., Sargent, D.J., Hancock, J. and Bassil, N.V. (2017) Genotyping-by-sequencing enables linkage mapping in three octoploid cultivated strawberry families. PeerJ, 5, e3731 10.7717/peerj.3731.

Wen, Z., Yao, L., Wan, R., Li, Z., Liu, C. and Wang, X. (2015) Ectopic Expression in Arabidopsis thaliana of an NB-ARC Encoding Putative Disease Resistance Gene from Wild Chinese Vitis pseudoreticulata Enhances Resistance to Phytopathogenic Fungi and Bacteria. Frontiers in Plant Science, 6, 1087

Witek, K., Jupe, F., Witek, A.I., Baker, D., Clark, M.D. and Jones, J.D.G. (2016) Accelerated cloning of a potato late blight–resistance gene using RenSeq and SMRT sequencing. Nature Biotechnology, 34, 656 10.1038/nbt.3540 https://www.nature.com/articles/nbt.3540#supplementary-information.

Xiao, S., Ellwood, S., Calis, O., Patrick, E., Li, T., Coleman, M. and Turner, J.G. (2001) Broad-Spectrum Mildew Resistance in Arabidopsis thaliana Mediated by RPW8. 10.1126/science.291.5501.118.

Xiong, J.-S., Zhu, H.-Y., Bai, Y.-B., Liu, H. and Cheng, Z.-M. (2018) RNA sequencing-based transcriptome analysis of mature strawberry fruit infected by necrotrophic fungal pathogen Botrytis cinerea. Physiological and Molecular Plant Pathology, 104, 77–85 https://doi.org/10.1016/j.pmpp.2018.08.005.

Yang, Z. (2007) PAML 4: phylogenetic analysis by maximum likelihood. Molecular biology and evolution, 24, 1586–1591

Zhong, Y. and Cheng, Z.-M.M. (2016) A unique RPW8-encoding class of genes that originated in early land plants and evolved through domain fission, fusion, and duplication. Scientific Reports, 6, 32923 doi:10.1038/srep32923.

Zhong, Y., Zhang, X. and Cheng, Z.-M. (2018) Lineage-specific duplications of NBS-LRR genes occurring before the divergence of six Fragaria species. BMC Genomics, 19, 128 10.1186/s12864-018-4521-4.

