## Supplementary Files for "Disease Resistance Genetics and Genomics in Octoploid Strawberry"

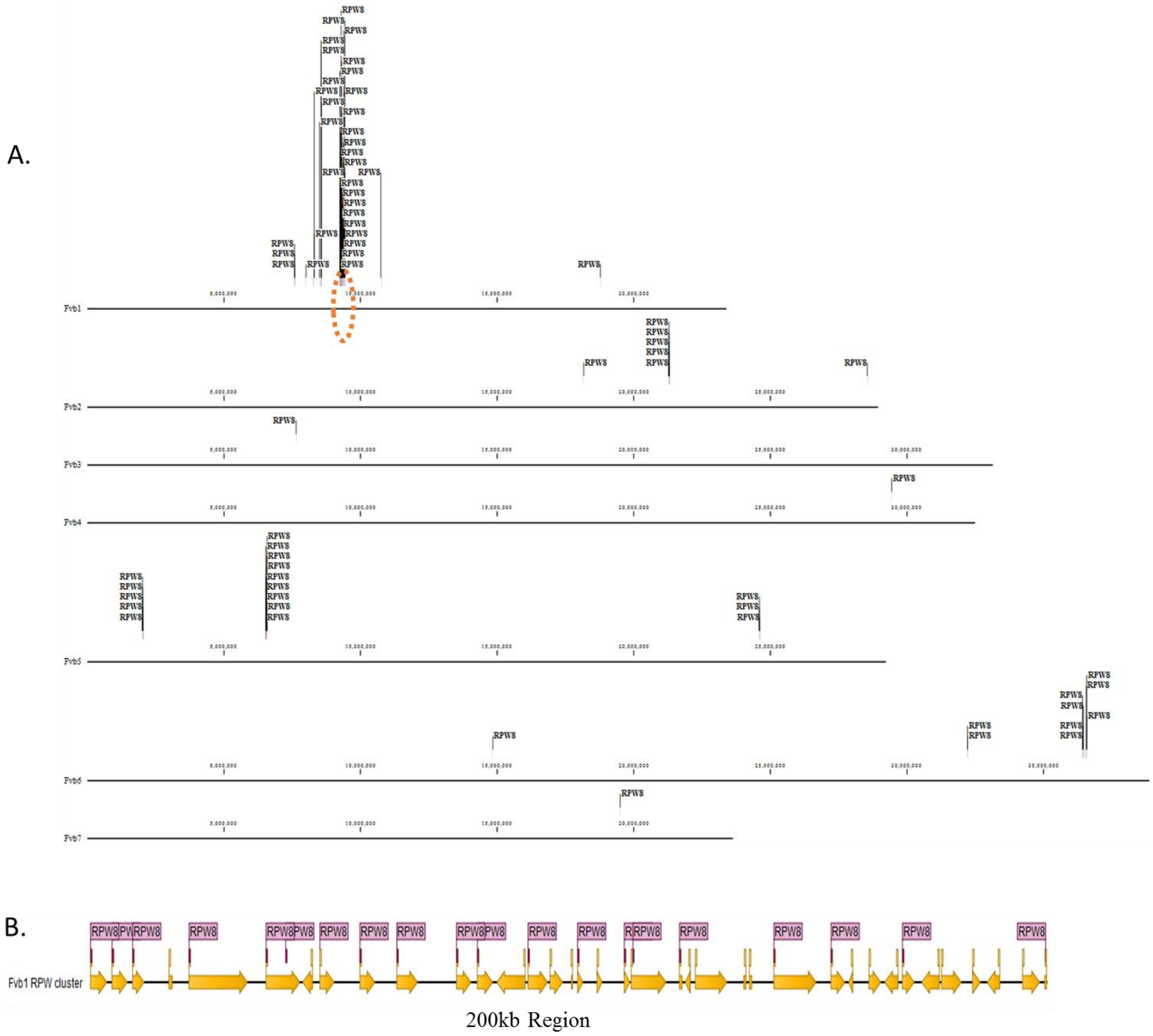


**Figure S1 Clustering of Similar R-gene Classes in F. vesca.** A. Most RPW8 domains in the F. vesca genome are arrayed in one of several genomic clusters, the most prominent of which is located on chromosome 1 (orange oval). B. A 200kb region of chromosome 1 (orange oval) contains a cluster of 18 tandemly-arrayed RPW8-containing R-genes.

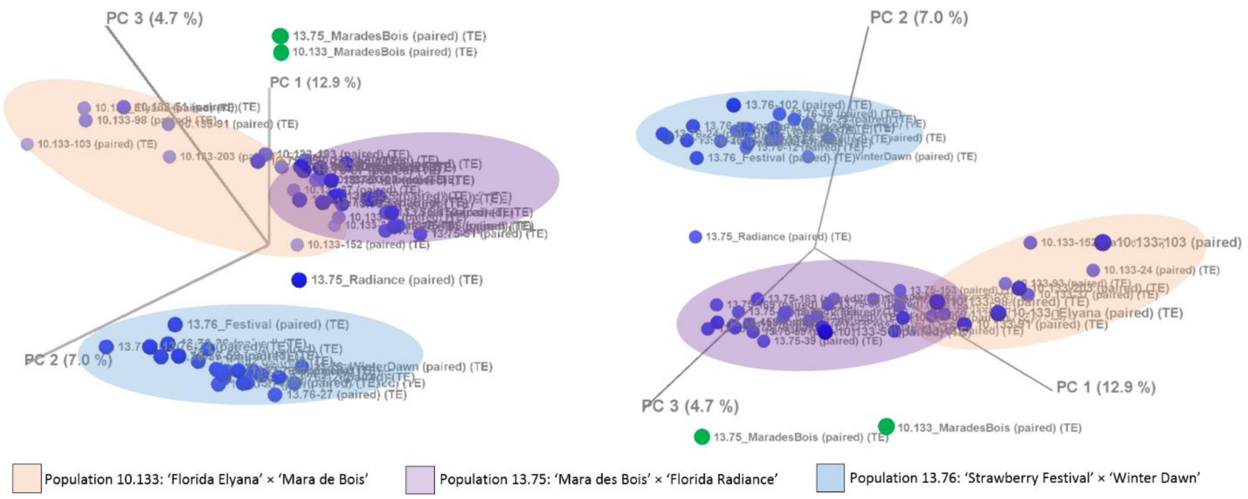


**Figure S2 Three-dimensional principle component analysis of RNAseq samples.** Populations "10.133" (orange), "13.75" (purple), and "13.76" (blue) are indicated. Two independent parental 'Mara des Bois' samples, collected with populations "10.133" and "13.75", are indicated with green dots. PC1, PC2, and PC3 represent 12.9%, 7.0% and 4.7% of total transcriptional variance, respectively. Two angles of the three-dimensional PCA show the minimal and maximal 2D distances between 'Mara des Bois' samples.

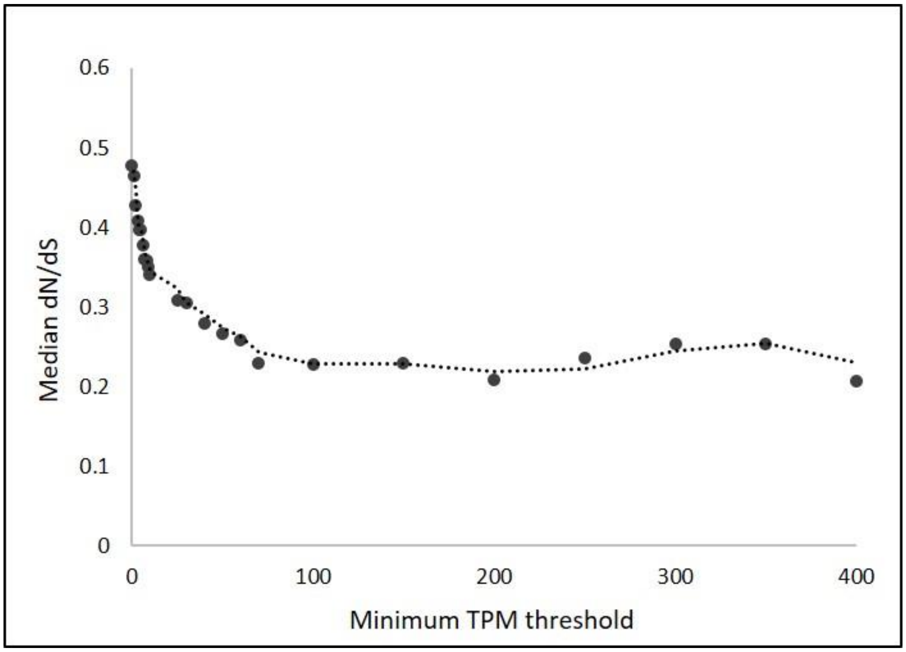


**Figure S3 R-gene *d*N/*d*S ratios compared with transcript abundance.** Low transcript accumulation across all tissues is associated with a higher proportion of non-synonymous mutation (Pearson's $r = -0.69$, $p < .0001$).

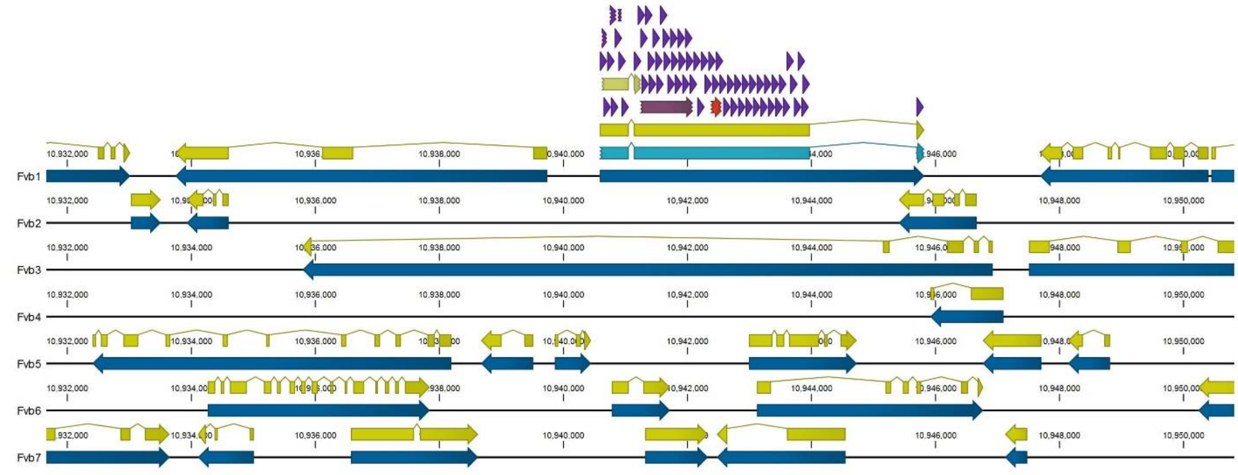


**Figure S4 Example RenSeq Probe Coverage of a TNL R-gene in the *F. vesca* Genome**. 120-mer RenSeq probes were designed against predicted R-genes (teal) in octoploid and diploid genomes. RenSeq capture probes (violet) do not span exon junctions, so they may be accurately applied to both gDNA and cDNA libraries. Annotations for Gene (blue), CDS (yellow), TIR (cream), NB-ARC (plum), and LRR (red) are shown.

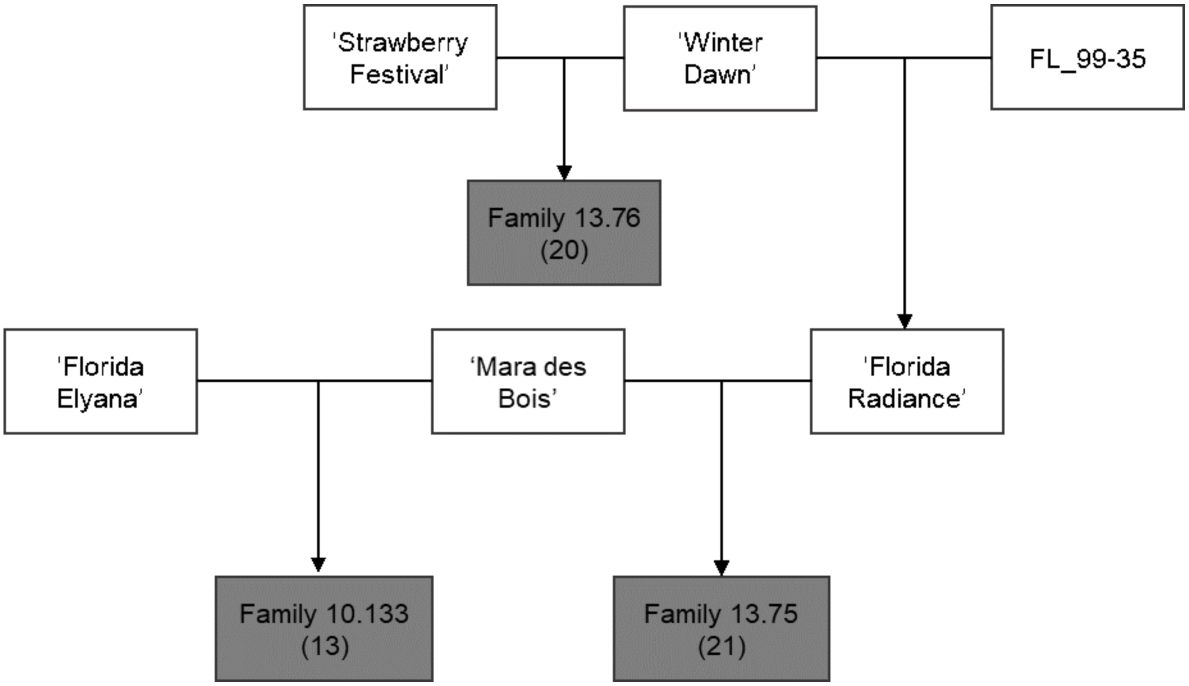


**Figure S5 Genetic Relatedness in the Three Segregating Octoploid Populations.** Pedigrees of three strawberry families used in RNAseq and eQTL analysis (dark gray), with number of sequenced progeny in parentheses.
